# ATAD3 megadalton complex in *Plasmodium falciparum* is essential for mitochondrial and cellular viability

**DOI:** 10.1101/2025.10.06.680775

**Authors:** Ijeoma C. Okoye, Ian M. Lamb, Yee-Wai Cheung, Joanne M. Morrisey, Swati Dass, Anurag Shukla, River S. Rell, Michael W. Mather, Yi-Wei Chang, Akhil B. Vaidya

## Abstract

Malaria remains an urgent threat to global health as the mortality and infection rates keep rising annually and our frontline antimalarials are becoming less effective due to the emergence and spread of resistance-conferring mutations. Although the mitochondrion of *P. falciparum* parasites is a validated drug target, there remain many uncharacterized mitochondrial proteins. The goal of this study was to investigate the essentiality and functions of a recently identified mitochondrial protein - PF3D7_0707400. Our results show that PF3D7_0707400 is an ATAD3A ortholog that is essential to parasite survival and is present in a megadalton complex that is critical for multiple mitochondrial processes such as mitochondrial RNA stability, membrane potential, ultrastructure, and protein import. This study is the first characterization, to our knowledge, of ATAD3A in unicellular organisms. ATAD3A has been previously studied in multicellular eukaryotes and has been implicated in several childhood mitochondrial diseases. Our findings here expand our knowledge on apicomplexan mitochondrial biology and our arsenal of potential antimalarial drug targets.

**Author Summary:** Each year, malaria is responsible for about 200 million infections and 600,000 deaths across the world. Thus, it constitutes a huge global health crisis. Increasing rates of antimalarial resistance necessitates the identification and characterization of novel parasitic proteins that can be exploited for the development of new antimalarial therapeutics. To this end, the mitochondrion of *P. falciparum* parasites has been studied as a validated target for effective antimalarials. However, much remains to be understood about critical mitochondrial processes and proteins that are essential for mitochondrial viability and parasite survival. Our study details the first characterization of an ATAD3 protein in a unicellular eukaryote, specifically in an apicomplexan parasite. The conservation of this protein in these deep-branching organisms highlights the importance of its biological functions, further emphasizing the significance of our study. By employing advanced molecular biology techniques, we show the presence of *Pf*ATAD3 in a giant molecular complex and its essentiality in asexual *P. falciparum* parasites. Conditional knockdown of *Pf*ATAD3 resulted in defects in critical mitochondrial processes such as mitochondrial RNA stability, mitochondrial membrane potential, and mitochondrial morphology. Divergence of *Pf*ATAD3 from the host allows for exploitation of this protein as a target for new antimalarials.

## Introduction

Malaria has plagued humanity ever since its dawn. Even today, more than half the world’s population is at risk of malaria infection. Hence, malaria remains a significant global health problem. Malaria is caused by unicellular apicomplexan parasites, *Plasmodium spp.,* transmitted by infected female *Anopheles* mosquitoes (1). Amongst the *Plasmodium spp*. that cause human malaria, *P. falciparum* is the species that causes the most fatalities (1, 2). The prevalence and incidence of malaria have increased in the last decade due to multiple factors including increased resistance to existing antimalarial therapies (3–10). Thus, there is a continued need to develop new and effective antimalarial drugs to counter the ever-present threat of drug resistance. The mitochondrion of *P. falciparum* is a validated antimalarial drug target (11, 12). Antimalarials already in use, such as atovaquone, target mitochondrial components (13). Each asexual *Plasmodium* parasite possesses a single mitochondrion that differs significantly from the human mitochondrion, having multiple divergent features and reduced metabolic function (14, 15). The *P. falciparum* mitochondrion houses about 30 copies of a 6 kb mitochondrial DNA element. This 6 kb DNA encodes mitochondrial ribosomal RNA fragments and three proteins that are components of Complexes III and IV of the *Plasmodial* electron transport chain – cytochrome *c* oxidase I and III (COX I/COXIII), and cytochrome *b* (Cyt *b*) (16–19). Aside from the three proteins made in the mitochondrion, there are more than 400 proteins that are encoded by the parasite’s nuclear genome but are imported into the mitochondrion to perform an array of critical functions including mitochondrial DNA replication/transcription, mitochondrial mRNA translation, ubiquinone biosynthesis, pyrimidine biosynthesis, and Fe-S cluster synthesis (12). However, several essential mitochondrial processes, such as tRNA import into the mitochondrion and mitochondrial ribosomal assembly, are still poorly understood. Therefore, much remains to explore and understand about mitochondrial biogenesis and functioning in *P. falciparum* parasites, with a view to identify potential targets for selective inhibition by new antimalarials.

Previously, through proximity biotinylation with mitochondrially targeted biotin ligase TurboID, we identified 122 putative mitochondrial proteins of *P. falciparum* parasites (20). One of these proteins was PF3D7_0707400, annotated as ATPase family AAA+ (<u>A</u>TPases <u>a</u>ssociated with diverse cellular <u>a</u>ctivities) domain containing protein 3 (*Pf*ATAD3). *Pf*ATAD3 is distantly related to a human protein, ATAD3A, which is a nuclearly encoded protein that has been demonstrated to have or modulate multiple mitochondrial functions including mitochondrial nucleoid stabilization, mitochondrial protein synthesis, mitochondria-endoplasmic reticulum inter-organellar interactions, and mitochondrial inner membrane protein scaffolding (21–31). Mutations in ATAD3 are implicated in many childhood mitochondrial diseases and are responsible for a variety of disorders in humans, ranging from neurological diseases to cancers (32–38). Until now, there has been no experimental characterization of this protein in *P. falciparum* or other apicomplexan parasites. This study represents the first characterization, to our knowledge, of ATAD3 in a unicellular system. Here, we demonstrate that *Pf*ATAD3 is essential for the growth of *P. falciparum* parasites and is present in a large multi-megadalton complex involved in critical mitochondrial processes.

## Results

### *Pf*ATAD3 is an ortholog of human ATAD3A and is found in the *P. falciparum* mitochondrion

Human ATAD3A is a 586-amino-acid protein classified as belonging to the AAA+ ATAD3 family of proteins with conserved ATAD3 and ATPase domains at its N and C termini, respectively. It has additional features such as a proline rich motif (PRM) and coiled-coil regions (CC1 and CC2) at its N-terminus, which extend into the cytoplasm of the cell, as well as transmembrane domains (TM1 and TM2), which span the inner and outer mitochondrial membranes (Fig 1A). *Pf*ATAD3 is a 663-amino-acid protein that possesses the conserved N-terminal ATAD3 and C-terminal AAA+ family ATPase domains, as well as potential TM helices, but lacks the PRM (Fig 1A). As AAA+ ATPases are known to exist as hexamers, we carried out a comparison of the predicted 3D structures of the hexameric human ATAD3A and *Pf*ATAD3 protein complexes, revealing a probable significant structural homology (Fig 1A). A pairwise global alignment between human ATAD3A and *Pf*ATAD3 showed 29.59% sequence identity between the two proteins (Fig 1B). We performed a phylogenetic analysis to determine the evolutionary relationships of ATAD3A in apicomplexan parasites and other eukaryotes (S1A Fig). Orthologs of *Pf*ATAD3 exist in other apicomplexan parasites and Myzozoans, including *Cryptosporidium parvum* which has no mitochondrial DNA but has a mitosome (a mitochondrion-derived organelle that does not produce ATP). Plants and algae also possess ATAD3A orthologs (S1A Fig). Interestingly, there are no ATAD3A orthologs in fungi. A multiple sequence alignment of the ATAD3 orthologs across apicomplexan parasites revealed a significant conservation in sequence similarity, including the Walker A, Walker B, and Arginine Finger domains that are crucial for ATPase activity and oligomerization (S1B Fig). *Plasmodium spp.* parasites tend to have distinctly long C-terminal extensions in addition to the conserved domains, indicating potential specific functions of ATAD3 in *Plasmodium* (S1B Fig).

**Figure 1.**
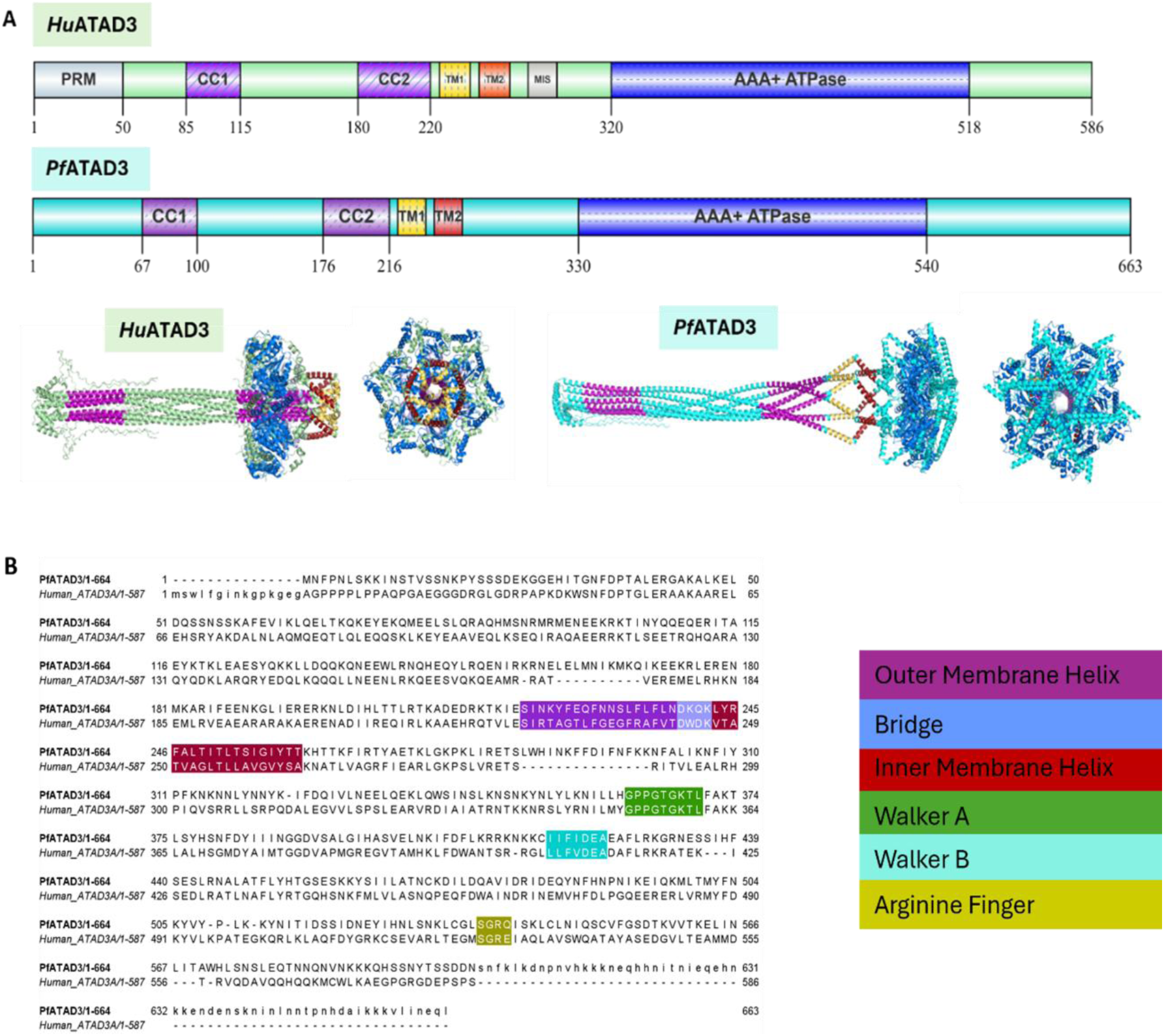
Bioinformatic analysis of *Pf*3D7_0707400. **(A)** Sequence and structural homology analysis of human ATAD3A (Above) and *Pf*3D7_0707400 (Below). Sideview orientations on the left side and top-down view on the right side. Domains are distinguished by colors: Coiled-coil domains in purple, Transmembrane 1 (TM1, Outer Membrane) domain in yellow, Transmembrane 2 (TM2, Inner Membrane) domain in red, ATPase domain in dark blue. **(B)** Pairwise sequence alignment between *Pf*3D7_0707400 and human ATAD3A indicating 29% sequence identity and a conservation of critical domains including the Walker A domain (highlighted in green), the Walker B domain (highlighted in light blue), and the conserved Arginine finger (highlighted in yellow).

Next, we sought to determine the localization of *Pf*ATAD3 through immunofluorescence microscopy. To facilitate this, we generated endogenously 3xHA-tagged parasites using CRISPR/Cas9 homology-directed recombination (S2A-D Fig). The transgene was engineered to conditionally express *Pf*ATAD3 under TetR-DOZI aptamer regulation. Immuno-electron microscopy of these transgenic *Pf*ATAD3-TetR-3xHA parasites demonstrated *Pf*ATAD3 localization to the mitochondrion of asexual *P. falciparum* parasites (Figure 2A, S3A Fig). Localization of *Pf*ATAD3 to the mitochondrion has been previously validated through an immunofluorescence assay with MitoTracker using ectopically expressed *Pf*ATAD3 (20). Another transgenic parasite line was generated in which *Pf*ATAD3-3xHA-TetR parasites also express an mScarlet fluorescent tag targeted to the mitochondrion via the leader sequence of the mitochondrial chaperone protein *Pf*HSP70-3 (39). Immunofluorescence assay using these *Pf*ATAD3-3xHA-mito-mScarlet parasites showed significant colocalization of *Pf*ATAD3 with the mitochondrion across the ring, trophozoite, and early schizont asexual stages of the parasite (Fig 2B).

**Figure 2.**
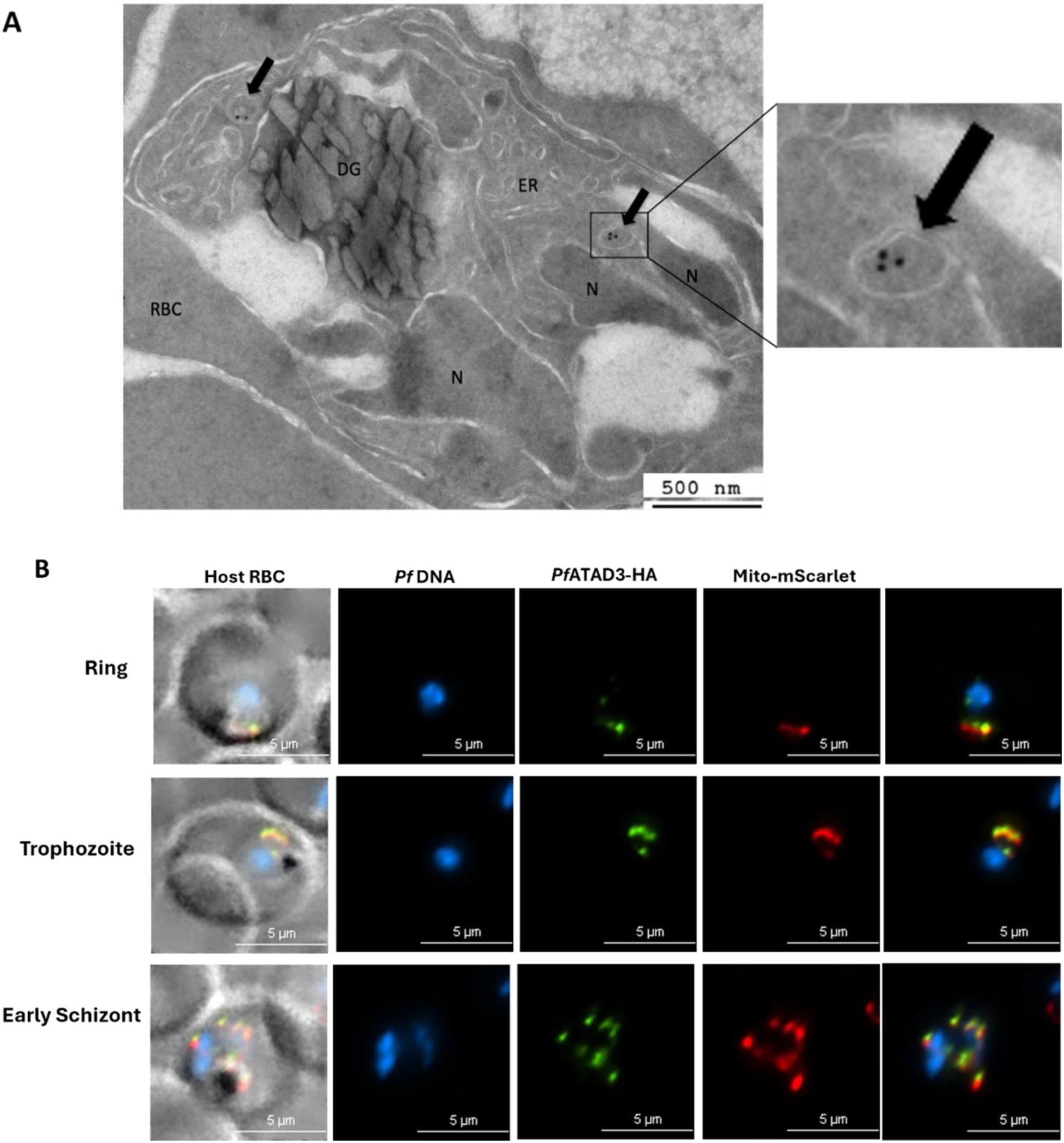
*Pf*ATAD3 is localized to the mitochondrion of asexual *P. falciparum* parasites. **(A)** Immuno-electron microscopy of *Pf*ATAD3-HA-TetR parasites demonstrating localization of *Pf*ATAD3 to the mitochondrion of asexual *P. falciparum* parasites using gold colloidal anti-mouse coated beads and mouse anti-HA antibodies. To the right is a zoomed-in view of *Pf*ATAD3 in the double-membraned mitochondrion. **(B)** Immunofluorescence Assay of *Pf*ATAD3-HA/mito-mScarlet parasites demonstrating localization to the mitochondrion across asexual stages of *P. falciparum* parasites.

### *Pf*ATAD3 is essential for the growth and development of asexual *P. falciparum* parasites

We generated a transgenic *Pf*ATAD3-3xHA-TetR parasite line also ectopically expressing *Pf*TOM22 (Translocator of Outer Mitochondrial Membrane 22) fused to a fluorescent mNeonGreen tag at its N-terminus to assess essentiality and functions of *Pf*ATAD3 in asexual stage *P. falciparum* parasites (S2E Fig, S3B Fig). Removal of aTc from the culture medium at ring stages resulted in ∼95% knockdown of *Pf*ATAD3 expression as early as 24 hours later (Fig 3A&B, S4 Fig). The deterioration of parasite morphology visualized by Giemsa staining (Fig 3C), together with measurement of the declining parasite growth rate via flow cytometry (Fig 3D), demonstrated that the parasites in the absence of *Pf*ATAD3 were in a state of growth arrest by 72 hours post aTc withdrawal and effectively died in the second intraerythrocytic asexual cycle. We further assessed the viability of *Pf*ATAD3(-) parasites by adding back aTc at different time points following the knockdown induced at ring stages. We observed that a significant proportion of 24 h and 48 h *Pf*ATAD3(-) parasites remained viable, as their growth resumed following aTc addition at these timepoints (Fig 3E). However, after 72 h and 96 h, a significant proportion of *Pf*ATAD3(-) parasites were no longer viable as they were unable to resume their growth after addback of aTc at these timepoints (Fig 3E). Thus, knockdown of *Pf*ATAD3 results in a growth arrest and eventual death of parasites within the second asexual cycle, indicating that *Pf*ATAD3 is essential for asexual *P. falciparum* parasite viability, growth and development.

**Figure 3.**
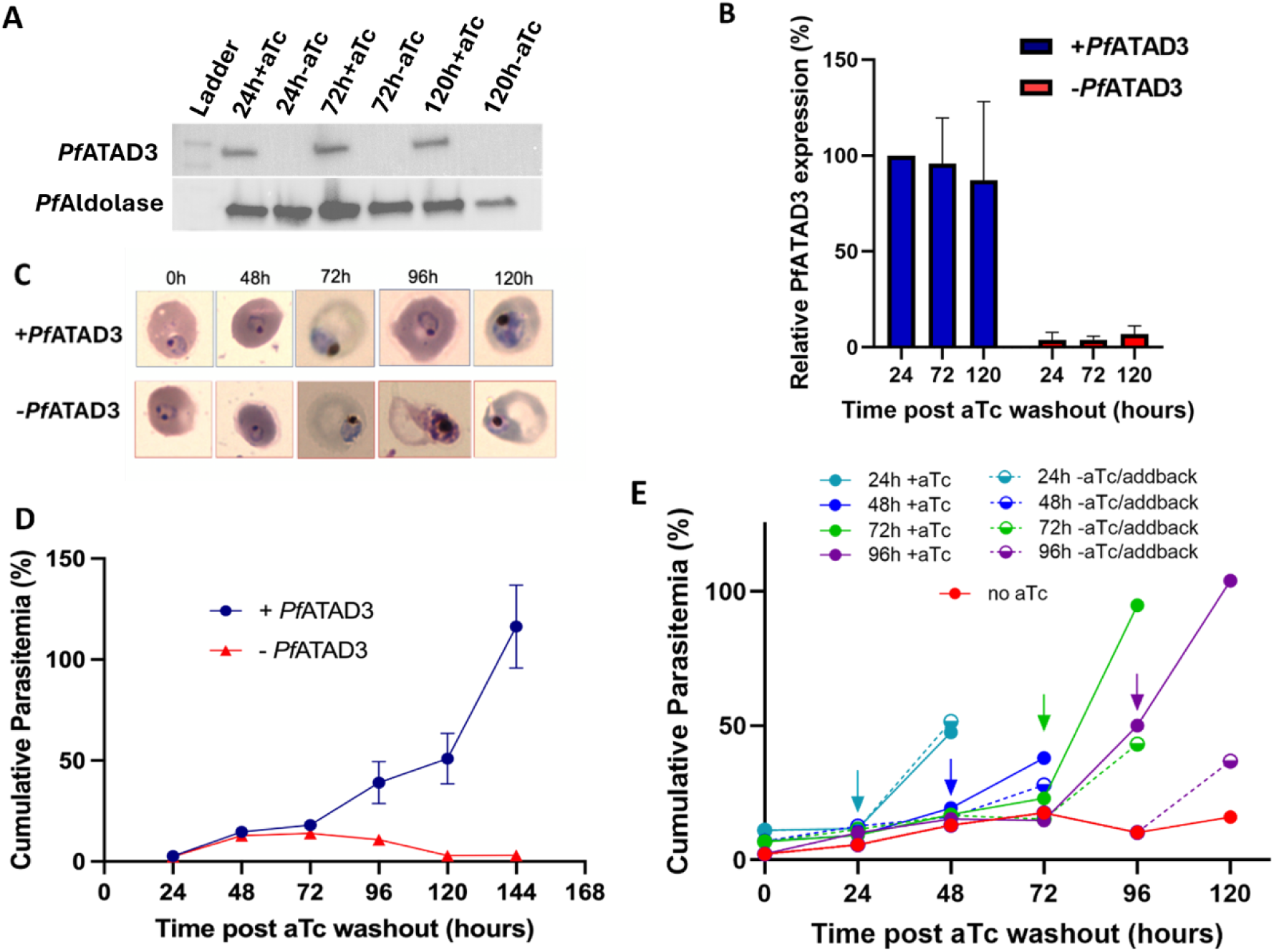
*Pf*ATAD3 is essential for asexual parasite survival and development. **(A)** Western blot and **(B)** Quantification of anhydrotetracycline (aTc)-mediated conditional regulation of *Pf*ATAD3 expression. *Pf*Aldolase was used as the loading control. The bar plot is an average of three independent biological replicates **(C)** Visualization of Giemsa-stained +/-*Pf*ATAD3 parasites by light microscopy indicating normal morphology and growth of *Pf*ATAD3(-) knockdown parasites for the first asexual cycle but the development of a growth arrest, aberrant morphology, and eventual death by the second asexual cycle. **(D)** Growth curve via flow cytometry of cumulative parasitemia for each condition (+/-*Pf*ATAD3) and each timepoint (every 24 hours for six days). 100,000 red blood cells were counted for each sample. The black curve represents *Pf*ATAD3(+) parasites while the red curve represents *Pf*ATAD3(-) knockdown parasites. The growth curve is an average of three independent biological replicates. **(E)** Growth curves demonstrating the loss of cell viability by 72h of *Pf*ATAD3 knockdown. Arrows indicate the timepoint that aTc was added back, i.e., 24h, 48h, 72h, and 96h. Solid lines represent +aTc control, dash lines represent the aTc addback condition, and the red line represents continuous –aTc knockdown condition.

### *Pf*ATAD3 is critical for mitochondrial function in asexual *P. falciparum* parasites

To assess the morphology and functionality of the mitochondrion in *Pf*ATAD3(-) parasites, we used the *Pf*ATAD3-3xHA-TetR*/Pf*TOM22 parasite line. *Pf*TOM22 imaging permitted visualization of the outer mitochondrial membrane morphology of the parasite, while staining of these parasites with MitoTracker Red dye enabled assessment of the mitochondrial membrane potential. Super-resolution live cell imaging of *Pf*ATAD3(+) parasites stained with MitoTracker showed colocalization with *Pf*TOM22 as expected due to a stable mitochondrial membrane potential (Fig 4A, S5A Fig). However, a loss of mitochondrial membrane potential was observed in *Pf*ATAD3(-) parasites 48 h after initiation of the knockdown, as revealed by the appearance of parasites with dispersed MitoTracker staining (Fig 4A, S5A Fig, Mov. S1 & S2). Meanwhile, outer mitochondrial membrane morphology in *Pf*ATAD3(-) parasites remained unaltered as indicated by *Pf*TOM22 organization in a normal tubular morphology after 48 hours of *Pf*ATAD3 knockdown (S5A Fig). Quantification of tubular and dispersed MitoTracker staining of parasites showed 60% of the *Pf*ATAD3(-) parasites had lost their mitochondrial membrane potential 48 h after *Pf*ATAD3 knockdown (Fig 4B). These observations suggest the importance of *Pf*ATAD3 for the maintenance of mitochondrial membrane potential, which contributes to mitochondrial physiology in asexual *P. falciparum* parasites. Notably, the continued retention of MitoTracker in the parasite cytoplasm suggests that the plasma membrane potential in the parasites remained intact at 48 h after knockdown initiation, even as the mitochondrial membrane potential was disrupted. This is consistent with our observation that the parasites were viable at 48 h following *Pf*ATAD3 knockdown in the aTc complementation growth assays.

**Figure 4.**
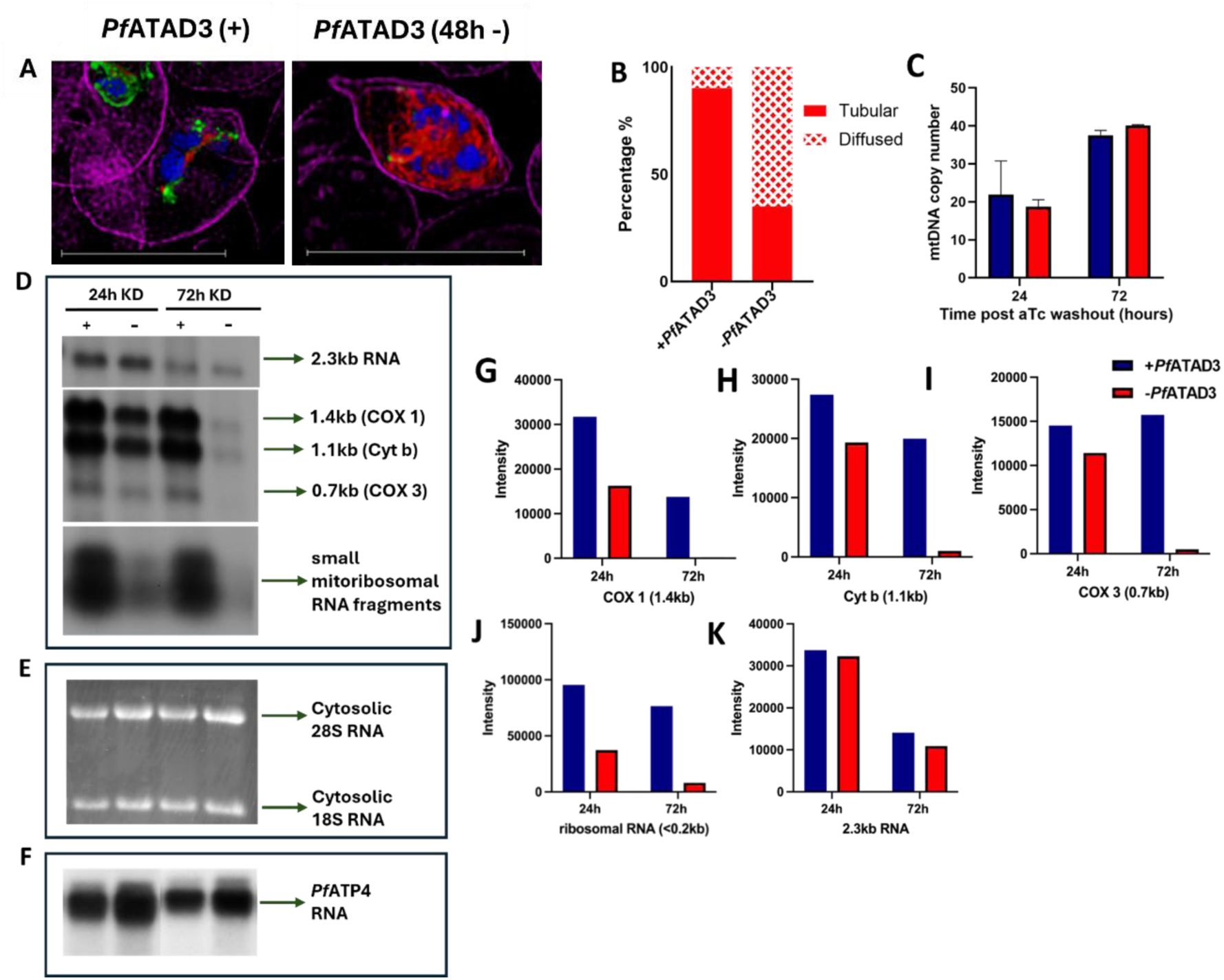
Genetic knockdown of *Pf*ATAD3 results in significant mitochondrial defects in asexual P. falciparum parasites. **(A)** Live Cell Scanning Confocal Super-resolution Microscopy of +/-*Pf*ATAD3 (48h) TetR/TOM22-mNG parasites [*Pf*DNA – DAPI (Blue); Outer mitochondrial membrane – *Pf*TOM22-mNG (Green); Mitochondrion – MitoTracker (Red); Host red blood cell – Wheat Germ Agglutinin (Purple). Scale bar is 10 µm **(B)** Quantification of cells with stable vs collapsed mitochondrial membrane potential based on super-resolution imaging in **(A)**. **(C)** Absolute quantification of cytochrome b copy number in +/- *Pf*ATAD3 parasites; average of two independent biological replicates **(D)** Northern Blot of mitochondrial messenger and ribosomal RNA transcripts in +/- *Pf*ATAD3 parasites. **(E)** Denaturing RNA gel electrophoresis demonstrating equivalent loading of total RNA isolated from +/- *Pf*ATAD3 samples as well as stability of cytosolic ribosomal RNA in +/- *Pf*ATAD3 parasites. **(F)** Northern Blot of nuclear-encoded *Pf*ATP4 messenger RNA in +/- *Pf*ATAD3 samples **(G-K)** Quantification of northern blot in **(D)** using FIJI software.

Next, we assessed the effect of *Pf*ATAD3 knockdown on the mitochondrial DNA and its transcription. Digital PCR was used for absolute quantification of the mitochondrial DNA, which showed no significant change in its copy number upon the knockdown of *Pf*ATAD3 (Fig 4C). Furthermore, we used northern blot analysis and denaturing RNA agarose gel electrophoresis to assess mitochondrial and cytoplasmic RNA levels respectively following *Pf*ATAD3 knockdown (Fig 4D and Fig 4E). At 24 h following *Pf*ATAD3 knockdown, the levels of mitochondrial mRNAs encoding Cyt b, COX I/III and of small mitochondrial rRNA transcripts were noticeably reduced (Fig 4D, Fig 4G-J). After 72 hours of *Pf*ATAD3 knockdown, minimal mitochondrial mRNA and rRNA transcripts were detected (Fig 4D, Fig 4G-J). However, cytosolic 18S and 28S RNAs were relatively stable irrespective of *Pf*ATAD3 knockdown (Fig 4E). Nuclear-encoded *Pf*ATP4 mRNA also remained relatively stable upon *Pf*ATAD3 knockdown (Fig 4F). Interestingly, a 2.3 kb RNA, which is likely an unprocessed precursor, remained mostly unchanged in its abundance during the knockdown (Fig 4K). Overall, these results show that *Pf*ATAD3 plays a crucial role in maintaining the stability and abundance of processed mitochondrial RNA transcripts in asexual *P. falciparum* parasites.

### *Pf*ATAD3 is important for functions beyond genesis/maintenance of the mitochondrial electron transport chain (mtETC)

The primary function of the mtETC in blood stage *P. falciparum* parasites is to support *de novo* pyrimidine biosynthesis due to the parasite’s inability to salvage pyrimidines (40). Mitochondrially localized dihydroorotate dehydrogenase (DHODH) is the fourth enzyme in this synthetic pathway and requires ubiquinone as the electron acceptor. The reduced ubiquinone needs to be re-oxidized to continue to serve as an electron acceptor for DHODH, a step catalyzed by Complex III of the mtETC. Treatment of parasites with Complex III inhibitors kills the parasites since ubiquinone regeneration is essential for parasite survival. However, supplementation with decylubiquinone can partially rescue the pyrimidine biosynthesis pathway and consequently the growth of the parasites even when Complex III is inhibited (41). To assess whether *Pf*ATAD3 has roles in addition to maintaining the mtETC activity, we induced knockdown of *Pf*ATAD3 in parasites supplemented with 25 uM or 50 uM decylubiquinone. In contrast to the *Pf*ATAD3(+) atovaquone-treated parasites whose growth defect was partially reversed upon supplementation with decylubiquinone, *Pf*ATAD3-knockdown parasites were not rescued by the addition of decylubiquinone (Fig 5A and 5B). Therefore, the functions of *Pf*ATAD3 appear to extend beyond maintaining stable mtETC activity.

**Figure 5.**
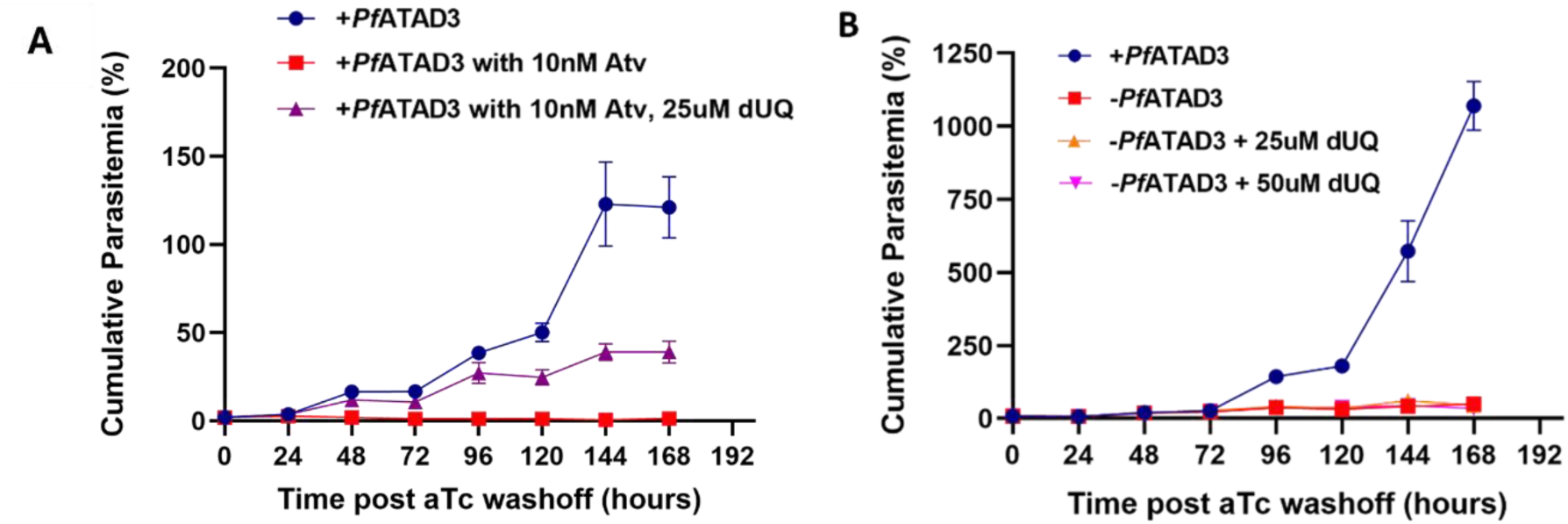
*Pf*ATAD3 plays critical roles in asexual *P. falciparum* parasites beyond mitochondrial electron transport chain activity. (A) Growth curve of cumulative parasitemia against time after drug (Atovaquone, Decylubiquinone) treatment. **(B)** Growth curve of cumulative parasitemia against time after PfATAD3 knockdown / Decylubiquinone treatment. The growth curve is an average of three independent biological replicates.

### *Pf*ATAD3 is present in a multi-megadalton complex in asexual *P. falciparum* parasites

AAA+ ATPases typically exist in a hexameric complex and are associated with additional proteins (42, 43). To determine if *Pf*ATAD3 exists within a complex, we proceeded to conduct native gel electrophoreses. Although a parasitophorous vacuolar membrane protein complex, *Pf*EXP2, migrated on a standard Blue-Native polyacrylamide gel (BN-PAGE) at the expected size of about 720 kilodalton, no band for *Pf*ATAD3 could be detected in the western blots of the BN-PAGE (S6A Fig). Treatment with minimal amounts of SDS separated the giant complex into its subcomplexes that could then be seen on a commercial BN-PAGE (S6A Fig). We reasoned that the native complex containing *Pf*ATAD3 was too large to enter the standard 4-12% BN-PAGE gel. Hence, we employed composite agarose-polyacrylamide native gel electrophoresis with a larger pore size than the standard BN-PAGE. On this gel, the *Pf*ATAD3 complex migrated at a position greater than 1 megadalton as judged from the migration pattern of *Pf*EXP2 complex (Fig 6A, S6B and S6C Figs).

**Figure 6.**
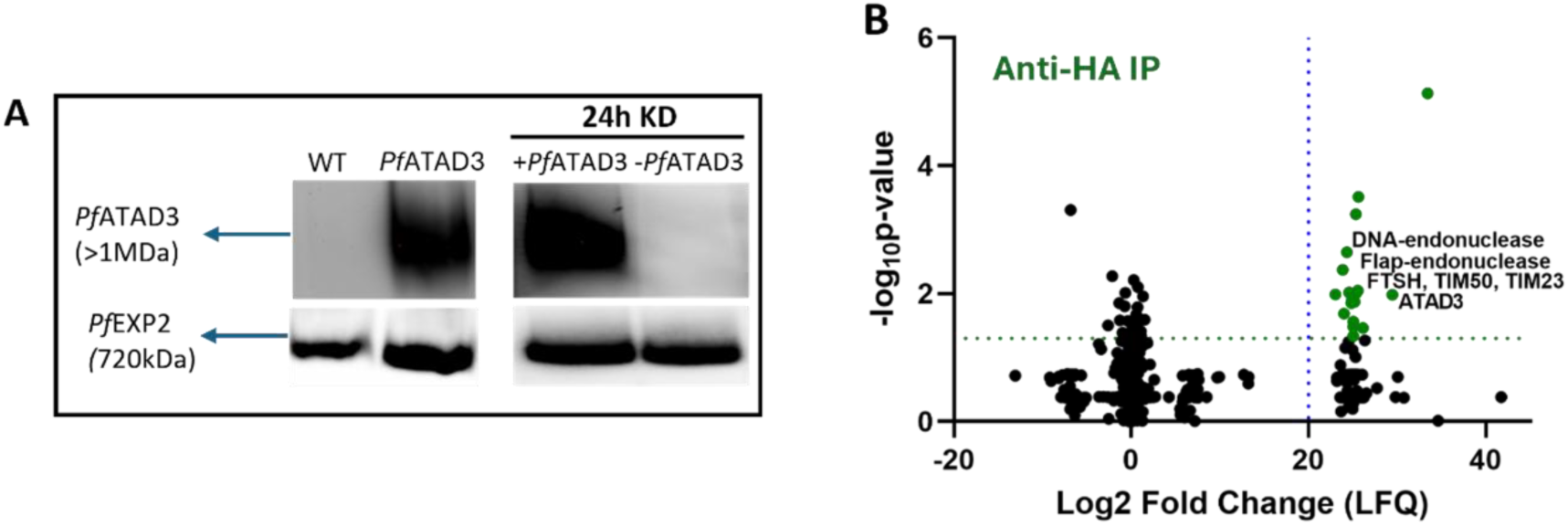
*Pf*ATAD3 exists in a multi mega-Dalton mitochondrial protein complex. **(A)** Large-pore composite agarose-polyacrylamide gel showing specific presence of >1MDa *Pf*ATAD3 complex in *Pf*ATAD3-HA (+) parasites **(B)** Volcano plot of fold change (log base 2) against the p-value illustrating proteins significantly enriched with *Pf*ATAD3.

To determine what proteins are present in this multi-megadalton complex, we carried out affinity isolation of *Pf*ATAD3 complex using anti-HA magnetic beads followed by mass spectrometry and proteomic analysis. As a negative control, HA affinity isolation was carried out with wildtype *P. falciparum* parasites lacking tagged proteins. Using a cutoff of a log_2_fold change enrichment of 20 and a p-value of 5% (p<0.05), we generated a list of proteins that were significantly enriched with tagged *Pf*ATAD3 relative to untagged controls (Fig 6B, S1 Table). Some of these enriched proteins have been assessed in published literature to be mitochondrially localized. As expected, *Pf*ATAD3 was the most enriched mitochondrial protein closely followed by inner membrane protein complexes such as another AAA+ ATPase protein PF3D7_1119600 (a putative *Pf*FTSH zinc metalloprotease) and Translocators of the Inner Mitochondrial Membrane, *Pf*TIM50 and *Pf*TIM23 (critical components of the mitochondrial import machinery). Mitochondrial DNA repair proteins such as DNA – (apurinic or apyrimidinic site) endonuclease (PF3D7_0305600) and flap endonuclease I (PF3D7_0408500) was also significantly enriched in the HA-pulldown. The significance of other proteins enriched with *Pf*ATAD3 remains to be investigated.

### Loss of *Pf*ATAD3 results in defects in the mitochondrial and cellular ultrastructure of asexual *P. falciparum* parasites

As ATAD3 proteins have been shown to be crucial for inner membrane proteins scaffolding as well as cristae formation (21, 44), we sought to assess the morphology of the mitochondrion as well as the cellular ultrastructure of parasites within the first (36 h) and second cycle (84 h) of *Pf*ATAD3 knockdown using transmission electron microscopy. Close examination of the mitochondrion in the parasites lacking *Pf*ATAD3 for 36 h reveals a marked increase in the width and area, as well as a reduction in the electron intensity in the mitochondrial matrix compared to their wildtype counterparts (Fig 7A-D). Furthermore, we observed significant aberrations in the intracellular membrane integrity and organization in the parasites without *Pf*ATAD3 for 84 h (Fig 7A-D). This finding is consistent with our observation of a loss of mitochondrial RNA and cellular viability at 72 h post *Pf*ATAD3 knockdown. Hence, *Pf*ATAD3 is important for maintaining proper mitochondrial morphology and, consequently, cellular viability in asexual *P. falciparum* parasites.

**Figure 7.**
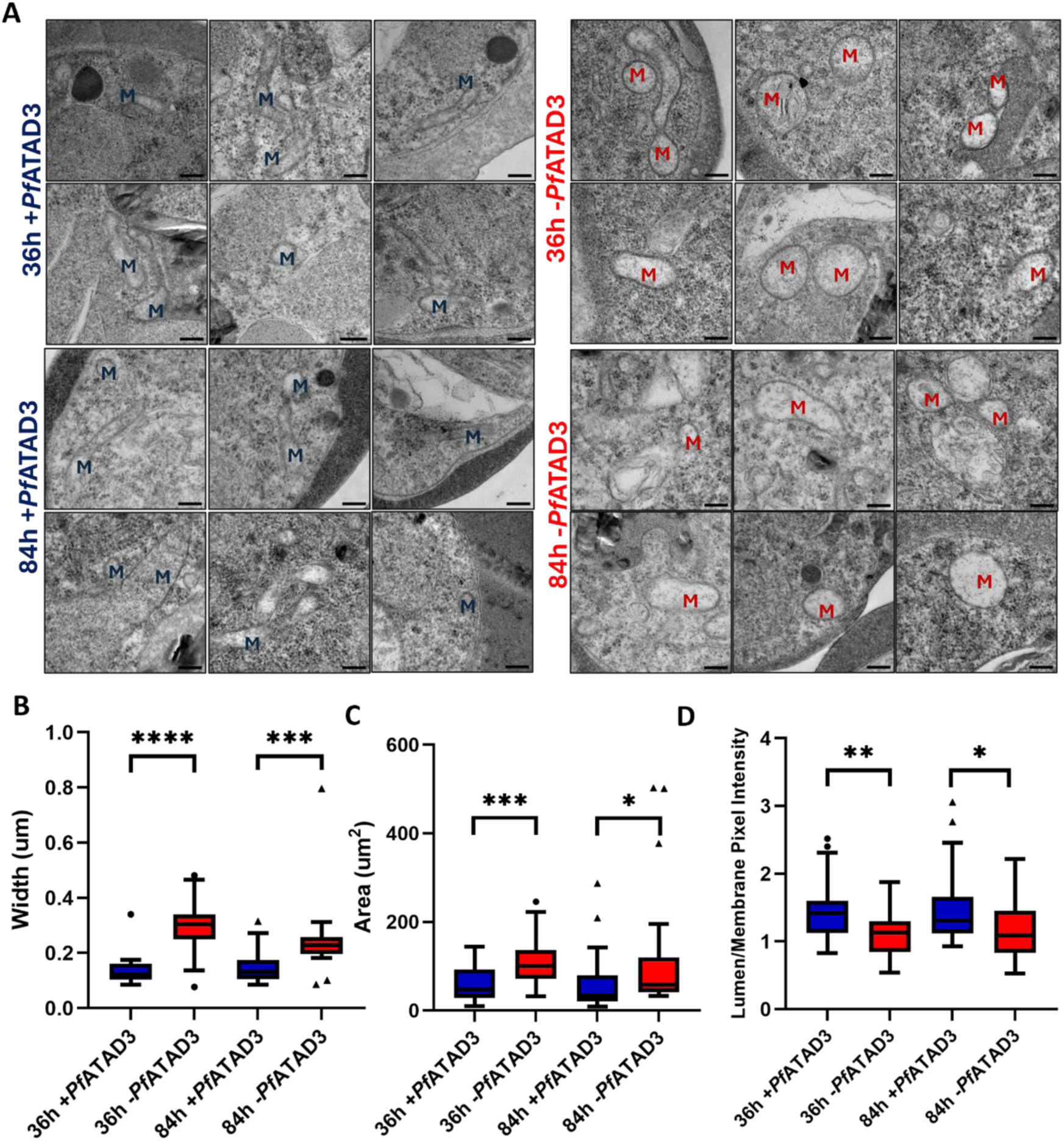
Impact of *Pf*ATAD3 knockdown on mitochondrial ultrastructure in asexual *P. falciparum* parasites. **(A)** Transmission electron micrographs of +*Pf*ATAD3 (control) parasites – left panels and –*Pf*ATAD3 (knockdown) parasites (right panels) at 36 h (top panels) and 84 h (bottom panels). Mitochondrion is marked as ‘M’. Scale bar is 200nm. **(B)** Quantification of the width (um), **(C)** area (um^2^), and **(D)** pixel intensity of the mitochondrion on FIJI. Statistical analyses were conducted via unpaired t-tests.

## Discussion

In this report we characterize ATAD3, a mitochondrial AAA+ ATPase, from the unicellular eukaryote *P. falciparum*. All previous studies on ATAD3 have been performed from multicellular eukaryotes (31). Indeed, ATAD3 was initially thought to be confined only to multicellular eukaryotes (24). Therefore, our investigation represents the first exploration of its apparent ortholog in a unicellular eukaryotic mitochondrion. In multicellular eukaryotes, ATAD3 has been associated with a myriad of functions such as mitochondrial inner membrane protein assembly and structure, mitochondrial DNA nucleoid dynamics, lipid metabolism, and cholesterol trafficking (21, 22, 25, 27, 30, 35, 36, 45). Its deletion results in embryonic lethality in worms, fruit flies, and mice (24, 44, 46, 47). In humans, multiple mutations in ATAD3 gene have been associated with neurological and developmental disorders including various forms of cancer (32, 34, 37, 48, 49). Interestingly, the ATAD3 gene is absent among the fungal lineage, yet it is present in much more evolutionarily distant organisms such as apicomplexan parasites. As is characteristic of all AAA+ ATPases, ATAD3 proteins have the conserved ATPase domain comprising of Walker A and B domains that allow for the binding and hydrolysis of ATP, respectively. The energy derived from ATP hydrolysis drives molecular processes attributed to AAA+ ATPases such as translocation of macromolecules, including nucleic acids and proteins, enabling DNA unwinding and replication, as well as protein unfolding and transport (33, 43, 50). Previous studies have described a fascinating topology of human ATAD3A as a membrane protein complex with its C-terminal ATPase domain localized in the mitochondrial matrix, its transmembrane domains I and II anchoring the complex in the inner and outer mitochondrial membranes, and its N-terminal coiled-coil domain extending into the cytoplasm to mediate interactions with other organelles such as the endoplasmic reticulum (ER) (24, 28). Although all the ATAD3 orthologs across humans and apicomplexan parasites have the conserved Walker A and Walker B ATPase motifs as well as the Arginine Finger, there are some features that are unique to human ATAD3A or *P. falciparum* ATAD3, respectively. Interestingly, human ATAD3A has a species-specific proline-rich motif (PRM) at its N-terminus that has been suggested to be involved in distinct functions such as mitochondria-ER inter-organellar interactions (22). In contrast, *P. falciparum* ATAD3 lacks such PRM but has a C-terminal extension of about 100 amino acids that may confer its own specific functions in *P. falciparum* parasites. Additionally, it is particularly notable that *Cryptosporidium* species, with a genome-less minimal mitochondrial remnant, also maintain ATAD3 genes. Conservation of ATAD3 over this vast evolutionary distance indicates essential but perhaps sometimes distinct functions that this mitochondrial protein may serve in different lineages of multicellular and unicellular eukaryotes.

Using a robust conditional knockdown system, we show here that *Pf*ATAD3 is an essential protein for development of asexual stages of *P. falciparum*. Parasites appear to remain in a growth arrested stage in the absence of *Pf*ATAD3 up to 72 h. Remarkably, we observed dramatic reduction of mitochondrially transcribed RNA as early as 24 h following the ATAD3 knockdown. Messenger RNAs as well as ribosomal RNAs encoded by the parasite mtDNA were barely detectable at 72 h following the knockdown. Cytoplasmic RNAs, however, did not seem to be affected. Interestingly, a 2.3 kb RNA moiety that is likely to be a precursor of mitochondrial mRNAs and rRNAs remained unchanged during the same period of *Pf*ATAD3 knockdown. Similarly, the copy number of mitochondrial DNA also remained unchanged during this period. We hypothesize that the absence of *Pf*ATAD3 affects processing and assembly of mitochondrial RNA transcripts. This would result in reduced production of mitochondrially encoded mtETC components and diminished respiration. Consequently, the arrested parasites underwent disruption of their mitochondrial membrane potential. This phenocopies previous observations with atovaquone treatment of asexual *P. falciparum* parasites where the parasites remain viable for a duration of time until eventual death despite an inhibited mtETC (51). However, the growth arrest does not appear to be due to a lack of ubiquinone regeneration (the main function of mtETC in the blood stage of the parasite to serve pyrimidine biosynthesis (40)), since addition of decyl-ubiquinone failed to rescue parasite growth in the absence of *Pf*ATAD3, in contrast to atovaquone-inhibited parasites, which are rescued by addition of decyl-ubiquinone (41). Thus, the absence of *Pf*ATAD3 appears to have a pleiotropic effect on mitochondrial functions in a manner that leads to parasite growth arrest and subsequent demise. This is consistent with existing literature describing altered mitochondrial functions including reduced expression of mtETC complexes and dysregulated respiratory activity upon loss or mutation of ATAD3 in multiple systems (21, 37, 45).

This defect of mitochondrial functions in *Pf*ATAD3 (-) parasites was also observed in tandem with defects in the mitochondrial ultrastructure of these parasites. Transmission electron microscopy revealed significant morphological changes in the mitochondrion of *Pf*ATAD3 (-) parasites. As early as 36 hours post *Pf*ATAD3 knockdown, the mitochondrion appeared swollen and with reduced internal electron density. This may be because of the collapsed mitochondrial membrane potential, as well as disruptions of mtETC complex assembly seen at 24 hours post *Pf*ATAD3 knockdown, which may be inhibiting critical processes such as protein import into the mitochondrion or assembly of essential mitochondrial inner membrane complexes. By 84 hours post *Pf*ATAD3 knockdown, the mitochondrion looked degenerated, coinciding with the data demonstrating loss of cellular and mitochondrial viability at this time. Changes in mitochondrial ultrastructure upon knockout or mutations of ATAD3 such as altered cristae structure, disrupted inner-outer mitochondrial membrane contact sites, electrolucent matrix areas, and overall degeneration have also been observed in mice and humans (35, 52, 53). Although asexual *P. falciparum* mitochondrion largely lacks cristae, our data shows that *Pf*ATAD3 still plays a critical role in maintaining normal mitochondrial morphology and ultrastructure in a unicellular eukaryotic organism.

To further understand the molecular functions of *Pf*ATAD3, we studied its native complex. Typically, AAA+ ATPase proteins oligomerize into hexameric complexes which can form pores (*46, 54, 55*). ATAD3 proteins are predicted to form hexameric complexes that have been suggested to traverse the inner and outer mitochondrial membranes (22, 42, 50, 54). Although *Pf*ATAD3 is predicted to also form similar hexameric complexes, we found it to be present in a much larger multi-megadalton complex that could only be displayed by composite large-pore native gel electrophoresis. This apparent size far exceeds six times the molecular mass of *Pf*ATAD3 that would be expected of a *Pf*ATAD3 hexamer (i.e., 468kDa), suggesting the presence of other molecules or complexes that are stably associated with *Pf*ATAD3. Proteomic analysis of immuno-isolated *Pf*ATAD3 complex revealed the possible identity of several associated proteins, including the components of the TIM23 complex (Translocase of Inner Membrane; TIM23, TIM17, TIM50), which mediate import of nuclearly-encoded inner mitochondrial membrane and matrix proteins critical for essential mitochondrial processes such as heme biosynthesis, Fe-S cluster biosynthesis, DNA/RNA replication, protein synthesis, cardiolipin synthesis, ubiquinone biosynthesis, and electron transport chain activity (55). Furthermore, another hexameric AAA+ ATPase, *Pf*FTSH, involved in protein homeostasis within the inner mitochondrial membrane was also pulled down with *Pf*ATAD3. *Pf*FTSH1 is annotated as an AAA+ ATP-dependent zinc metalloprotease, mediating mitochondrial protein processing and mitochondrial protein degradation. Experimentally, *Pf*FTSH1 has been shown to also localize to the mitochondrion of asexual parasites and oligomerize to form a hexameric complex >700 kDa, characteristic of AAA+ proteins (56). Higher order complexes were also observed suggesting the interaction of *Pf*FTSH1 with other proteins, which until now have not been identified (56). Our pulldown experiments also revealed interaction of *Pf*ATAD3 with DNA repair enzymes, *Pf*Ape1 and *Pf*Fen1; thereby, demonstrating a role of *Pf*ATAD3 in mitochondrial genome maintenance, which may impact abundance or expression of stable mitochondrial RNA transcripts. The specificity of these associations is emphasized by the lack of other integral mitochondrial membrane proteins such as cytochrome *b*, COX I/III, and mitoribosomal proteins in the pulldown. The interaction of *Pf*ATAD3 with specific inner membrane and matrix proteins indicate a likely role in various critical mitochondrial processes such as mitochondrial protein import, mitochondrial nucleoid stabilization, mitochondrial membrane potential, and mitochondrial ultrastructure. Other proteins annotated to localize in the endoplasmic reticulum and symbiont-containing vacuoles such as shewanella-like protein phosphatase and syntaxin respectively are also present in the *Pf*ATAD3-HA pulldown, indicating possible roles of *Pf*ATAD3 beyond the aforementioned mitochondrial functions.

Limitations of this study include evidence for a direct impact of *Pf*ATAD3 knockdown on import or localization of the nuclearly-encoded proteins or tRNAs into the mitochondrion. Isolation of mitochondria from parasites to directly and quantitatively assess impact of *Pf*ATAD3 knockdown on import and localization of mitochondrial proteins in *P. falciparum* is notoriously challenging due to contamination by hemozoin, the sticky crystalline by-product of hemoglobin degradation. We are also working on conducting a mutagenesis analysis to elucidate the essentiality and functions of distinct conserved domains in *Pf*ATAD3. Structural determination of *Pf*ATAD3 is also underway. Findings from structural and mutagenesis analysis of *Pf*ATAD3 would provide us with more details on the key residues and domains of *Pf*ATAD3 that can be exploited for rational structure-based drug design.

In conclusion, as is characteristic of AAA+ ATPases, *Pf*ATAD3 is involved in many important biological processes that are critical for mitochondrial biogenesis and functioning in asexual *P. falciparum* parasites. This paper provides the foundation for more in-depth and expansive analyses on the role of AAA+ ATPases, specifically ATAD3 proteins, in apicomplexan parasites and exploration of *Pf*ATAD3 as a potential target for novel antimalarial therapeutics.

## Materials and Methods

### Plasmid Construction

To generate a line in which *Pf*ATAD3A was endogenously tagged and expressed only in the presence of aTc, a gene containing homology regions 3’ and 5’ to the PF3D7_0707400 CRISPR/Cas9 induced double break site was synthesized by Genewiz (Azenta Life Sciences) (Fig. S1A). This gene was digested with AflII and BstEII and ligated into the pMGBKRML-*Pf*ATAD3-HA plasmid which contained the TetR and aptamer sequences (Fig. S1B and S1C). Prior to transfection, pMGBKRML-*Pf*ATAD3-HA vector was linearized by EcoRV digestion. For guide RNA construction, the pMKCas9 was linearized by EcoRI digestion and gel purified (Fig. S1D). Guide RNA inserts were selected using analysis from the Eukaryotic Pathogen CRISPR guide RNA design tool (RRID: SCR_018297). Guide RNAs were cloned into the EcoRI-linearized pMKCas9 vector by DNA assembly reaction (NEBuilder 59 HiFi DNA Assembly Master Mix, New England Biolabs, Inc.). The PF3D7_0707400gRNA1/2 primer sequences used in this reaction are listed in S2 Table.

### Parasite Lines, Culture, and Transfections

NF54 *attB P. falciparum* parasites were used in this study. Asexual *P. falciparum* parasites were cultured in human O^+^ red blood cells with RPMI 1640 medium supplemented with 0.5% w/v Albumax II (Fischer Scientific), sodium bicarbonate (2.1 g/L, Corning by ThermoFisher Scientific), HEPES (15 mM, ThermoFisher), hypoxanthine (10 mg/L, Fisher Scientific), and gentamycin (50 mg/L, VWR). To generate the *Pf*ATAD3-TetR-3xHA parasite line and the *Pf*ATAD3-TetR-3xHA/TOM22-mNeonGreen parasite line, wildtype NF54 parasites and *Pf*NF54-attB-TOM22-strepmNeongreen parasites generated as previously described (57) were transfected with 40 µg of linearized pMG75BB[3HA-10spApt-noP]-*Pf*ATAD3A plasmid as well as 20 µg of each gRNA plasmid. Transfections were performed on cultures containing 5-6% rings by electroporation via a Bio-Rad gene pulser (0.31 kV, 960 μFD). After electroporation, parasites were cultured under standard conditions for 48 hours before the addition of blasticidin (1.25 μg/mL, InvivoGen). The *Pf*ATAD3-TetR-3xHA parasites were grown in RPMI media containing 1.25 μg/mL blasticidin and 250 nM aTc while the *Pf*ATAD3-TetR-3xHA/TOM22-mNeonGreen were grown in RPMI media containing 1.25 μg/mL BSD, 250nM aTc, and 5 nM WR. To generate the parasite line where *Pf*ATAD3 was conditionally expressed and mScarlet was targeted to the mitochondrion, the PMG75-*Pf*ATAD3-tetRDOZI construct described above was transfected into the *Pf*NF54/iGPglmS line as previously described (58). The newly established *Pf*NF54iGP-*Pf*ATAD3 line was then transfected with a plasmid integrating the mitochondrial targeting sequence of Hsp70-3 fused to mScarlet into the SIL7 locus as previously described (39). This *Pf*ATAD3-3xHA-mitomScarlet line was maintained in RPMI media containing 2.5 mM glucosamine, 1.25 μg/ml blasticidin, and 250 nM aTc.

### Immunofluorescence Assay

50-100 µL of packed parasite cultures were pre-labeled with 60 nM MitoTracker Red CMXRos (Life Technologies by ThermoFisher Scientific) for 30 minutes at 37 °C. The samples were washed three times with 1X PBS. The cells were fixed with 4% v/v paraformaldehyde and 0.0075% v/v glutaraldehyde in 1X PBS for an hour at 37 °C with agitation and incubated overnight at 4 °C. The fixed cells were washed with PBS, permeabilized with 0.25% TritonX-100 for 10 minutes, and reduced with 0.1 mg/mL NaBH_4_ for 5 minutes at room temperature. The cells were blocked with 3% w/v bovine serum albumin (BSA) for 1 hour, incubated in primary mouse anti-HA antibodies 1:300 in BSA/PBS;(sc-7392 Santa Cruz Biotechnology) overnight at 4 °C, washed 3X in PBS and incubated in secondary Alexa Fluor 488 conjugated anti-mouse IgG 1:300 in BSA/PBS; (A32723 Invitrogen) overnight at 4 °C. Cells were also stained with DAPI. The cells were washed and resuspended in equilibration buffer and antifade (S2828 ThermoFisher Scientific) to preserve the samples and prevent photobleaching. 3 µL of each sample was mounted on a slide and imaged using a conventional inverted Nikon-Ti fluorescent microscope.

### Immuno-Electron Microscopy

NF54 *Pf*ATAD3-3xHA parasites were synchronized with alanine (0.3 M in 10 mM HEPES, pH 7.6) and enriched by a MACS Cell Separation Column (MiltenyiBiotec) at the trophozoite stage. 25-50 µL of the enriched trophozoites were fixed with 1 mL of 2% paraformaldehyde, 2.5% glutaraldehyde, and 100 mM sodium cacodylate for 1 hour on a rotator at room temperature. The cells were washed in sodium cacodylate buffer, embedded in 10% gelatin, and infiltrated overnight with 2.3 M sucrose and 20% polyvinyl pyrrolidone in PIPES/MgCl_2_ at 4 °C. The samples were then trimmed, frozen in liquid nitrogen, and sectioned using a Leica Ultracut UCT7 cryo-ultramicrotome. 50 nm ultrathin sections were then blocked with 5% fetal bovine serum and 5% normal goat serum for 30 minutes followed by incubation with primary mouse anti-HA antibodies (sc-7392, Santa Cruz Biotechnology) for 1 hour at room temperature. After washing in blocking buffer, the sections were incubated with 18 nm colloidal gold-conjugated secondary goat anti-mouse IgG (H+L) antibodies for 1 hour.

The sections were stained with 0.3% uranyl acetate and 2% methylcellulose. The stained sections were viewed on a JOEL 1200 EX transmission electron microscope with an AMT 8-megapixel digital camera and AMT Image Capture Engine V602 software (Advanced Microscopy Techniques). All labeling experiments were done in parallel with controls lacking the primary antibody. These controls were consistently negative at the concentration of colloidal gold-conjugated secondary antibodies used in these experiments. Immuno-electron microscopy was conducted at the Molecular Microbiology Imaging Facility at Washington University, St. Louis, MO.

### Parasite Growth Assay

The parasite culture was synchronized with 0.3 M alanine for two cycles to ensure all parasites were at the ring stage. After 48 hours, the parasites were washed three times with regular media to remove the aTc. The culture was split into two flasks where one was maintained with WR, aTc, blasticidin media serving as a control culture while the other was maintained with only WR99210 and blasticidin media serving as the knockdown culture. A 30-50 µL cell pellet was collected from each flask every 24 hours, fixed with 4% paraformaldehyde and 0.0075% glutaraldehyde for 1 hour at 37 °C and overnight at 4 °C. The fixed cells were washed with 1X PBS, stained with SYBR Green at a 1:10000 dilution, and then resuspended in filtered deionized water at a 1:100 pellet to water ratio. The samples were then assessed for parasitemia via flow cytometry. Thin blood smears were also made from each flask 24 hours after washout of aTc and stained with GIEMSA to manually calculate the parasitemia but also observe any morphological changes in the parasites after *Pf*ATAD3 knockdown.

### Western Blotting

Parasite cultures were lysed with 0.14% w/v saponin in 1X PBS supplemented with 1X protease inhibitor cocktail (P8215, MilliporeSigma). The parasite pellet was resuspended in 1X SDS/sample buffer and heated at 65 °C for five minutes. Samples were centrifuged at 12,000rpm for five minutes and the supernatant from each sample was run on a 10% BioRad Tris-glycine polyacrylamide gel. The proteins were electroblotted onto a methanol activated PVDF membrane. The membrane was blocked in 5% fat-free milk for 1.5 hours and washed with 1X TBS/Tween (TBST) before being incubated in primary mouse anti-HA antibodies (sc-7392, Santa Cruz Biotechnology) at 1:10000 in 1% fat-free milk overnight. The membrane was washed four times with TBST for 15 minutes each and incubated in a horseradish peroxidase-conjugated (HRP) secondary goat anti-mouse antibody (A16078, ThermoFisher Scientific) overnight at a 1:10000 dilution in 2% fat-free milk. The membrane was developed with SuperSignal Dura substrate (Pierce) and imaged. As a loading control, the blots were later re-blocked, washed, and probed with a rabbit anti-*Pf*Aldolase at a 1:30000 dilution followed by a HRP secondary mouse anti-rabbit antibody at 1:30000. All other steps followed the standard protocol.

### Live Cell Imaging

Glass bottom culture dishes (35 mm) were coated with 0.1% poly-L-lysine overnight and washed 3 times with PBS. Parasite culture (250 µL at 2.5% hematocrit) was added to the dishes and incubated for 30 min at 37 °C to allow attachment of parasites to the culture dish. Culture dishes were washed 3 times with PBS followed by addition of phenol red-free RPMI culture medium into which MitoTracker at 61 nM final concentration and Hoechst at 10 µg/mL final concentration were added and incubated for 30 and 20 minutes at 37 °C respectively to stain the mitochondrion and DNA respectively. Imaging was done using the Nikon Ti fluorescent microscope with the stage heater set to 37 °C. Hoechst was visualized using a DAPI (49,6-diamidino-2-phenylindole) filter set, mNeonGreen with (fluorescein isothiocyanate) FITC, and MitoTracker was visualized using tetramethyl rhodamine isocyanate (TRITC) filter set.

### DNA and RNA Isolation

Red blood cells were lysed using 0.14% w/v Saponin to isolate parasites and washed three times with 1X PBS. DNA was isolated according to the DNA isolation from blood protocol in the QIAGEN kit. For RNA isolation, the parasite pellet was resuspended in TriZol and then RNA was extracted using the phenol-chloroform phase separation method. Extracted RNA was run on a 1% denaturing RNA agarose gel to assess RNA integrity and the concentration/purity of RNA was determined via Nanodrop.

### Digital PCR

Custom QuantStudio Absolute Q DNA Digital PCR Assays were developed to determine the absolute copy number of Cytochrome b and GAPDH as measures of mitochondrial DNA and nuclear DNA copy number at 24 hours and 72 hours post *Pf*ATAD3 knockdown. Each assay consists of custom designed primers that are linked to FAM and VIC fluorescent probes to allow detection of Cyt b and GAPDH amplicons respectively (S3 Table). A 9 µL reaction mix was prepared for each sample, consisting of 1.8ul of Absolute Q DNA Digital PCR Master Mix (5X), 0.45ul of Cyt b – FAM Digital PCR Assay (20X), 0.45ul of GAPDH – VIC Digital PCR Assay (20X), 1 µL of 0.1ng/µL of genomic DNA from +/- *Pf*ATAD3-HA/TOM22-mNG parasites, and 5.3ul of nuclease-free water. Each 9 µL reaction was loaded to the bottom of the well and 15 µL of the Absolute Q Isolation buffer was loaded on top in the same well. All columns of the micro-array plate (MAP16) were sealed and loaded onto the Quant Studio Digital PCR machine. PCR thermal parameters were set to preheat to 96 °C for 10 minutes, denature at 95 °C for 3 seconds, anneal and extend at 60 °C for 30 seconds (20 cycles). Both FAM and VIC channel dyes were selected to detect copies of Cytochrome b and GAPDH genes respectively. Genomic DNA from wildtype NF54 parasites was used as a positive control and nuclease-free water was used as a negative control.

### Northern Blotting

20 µg of total RNA was mixed with RNA loading dye, 6% formaldehyde, 50% formamide, and 1X MOPS electrophoresis buffer. The samples were incubated in a 68 °C water bath for 5 minutes. The samples were loaded on a 1.2% agarose-2.2 M formaldehyde gel in 1X MOPS/DEPC buffer and run at 35 V at room temperature for 18-20 hours. One half of the gel was stained with ethidium bromide and visualized using a UV transilluminator to assess the presence of intact ribosomal RNA bands. The remaining gel half was rinsed 4x with DEPC-treated water to remove formaldehyde. RNA was transferred via capillary action 18-20 hours using 10xSSC onto a Genescreen PLUS nylon membrane, prewet with DEPC water and equilibrated with 10XSSC. After transfer the membrane was rinsed briefly in 2XSSC, air dried, and baked at 80 C in a vacuum oven for 2hrs. Ethidium Bromide staining and visualization of the gel post transfer indicated successful transfer of nucleic acids to the membrane as indicated by the lack of nucleic acids in the gel. The baked membrane was briefly rinsed in 2X SSPE buffer and prehybridized in 0.05 mL/cm^2^ hybridization solution containing 2X SSPE, 50% deionized formamide, 1% SDS, 5X Denhardts, and 10% Dextran Sulfate in a sealed bag for 2-4 hours at 42 °C with gentle agitation. The blots were probed with gel purified 6kb PCR of *P. falciparum* mitochondrial DNA labeled with Easytides dATP a-p32 (Perkin Elmer BLU512Z 6000Ci/mmol) using the Random Primer labelling method. The labelled probes were purified using an Illustra ProbeQuant G-50 micro column (GE Healthcare 28-0934-28) and labelling confirmed using Cerenkov counting in a Packard Tricarb LSC. Sheared calf thymus DNA at 100 µg/mL and probe at 1x10^6^cpm/mL were heated together for 5 minutes at 95 °C and placed on ice for 15 minutes. The probe was added to the blot in the bag. The bag was resealed and incubated at 42 °C for 18-24 hours with gentle agitation. The blot was removed from the bag and rinsed with 100-200 mL 2xSSC buffer at room temperature for 30 minutes, washed twice with 200 mL 2XSSC, 1% SDS at 50 °C for 30 minutes each, and washed with 0.2XSSC at room temperature for 30 minutes. The signal on the blot was checked after each wash using a Ludlum model 3 Geiger counter. The blot was drained, placed on a damp Whatman filter paper, covered with plastic wrap, exposed to X-ray film for 24-72 hours at -80 °C, and developed.

### Blue-Native Page Electrophoresis

Saponin isolated Parasite pellets were solubilized with solubilization buffer (200 mM 6-aminocaprioc acid, 50 mM Bis-Tris, pH 7, 1 mM EDTA, 2% digitonin, protease inhibitor cocktail 1:1000 Sigma P1812,1 mM AEBSF) overnight at 4 °C. The samples were centrifuged at 12000 rpm for 10 minutes and the supernatants were collected. Supernatants were run on a 4-12% gradient BN-PAGE gel as described in manufacturer’s standard protocol (Invitrogen). The ladder was fixed with 40% methanol/10% acetic acid for 30 minutes, stained with colloidal Coomassie, and then destained with 8% acetic acid. The proteins in the gel were transferred to a PVDF membrane and visualized by the methods described previously.

### Large Pore Composite Agarose-Polyacrylamide Gel

#### Electrophoresis

6 mL of deionized water, 1 mL of 30% bis/acrylamide, 1 mL of 20X Native Buffer (Invitrogen), 1.6 mL of 30% glycerol, 200 µL of 1M MgCl_2_, 200 µL of 10% ammonium persulfate (APS), 10 mL of 1% melted agarose in deionized water, and 20 µL of TEMED were mixed in order. The gel was cast using BioRad minigel 1.5 mm glass plates with a 6-well 1.5 mm comb. After solidifying for 1 hour at room temperature, the gel was submerged in 1X Native Running buffer and stored at 4 °C overnight. Samples were prepared in 1X Native Sample buffer (from Invitrogen NativePAGE Sample Prep Kit) / 5% G-250 and loaded in the wells. The gel was run at 140 V for 30 minutes and then at 100 V until the loading dye ran off. The gel was submerged in 1X SDS running buffer for 3 minutes before transferring proteins onto a PVDF membrane and visualizing as described previously.

#### Immunoprecipitation and Proteomic Analysis

NF54 wildtype parasites and *Pf*ATAD3-HA/TOM22-mNG parasites were cultured in large volume in triplicate and then lysed with 0.14% Saponin/1x Protease Inhibitor Cocktail. Parasite pellet was solubilized with 2% digitonin, protease inhibitor cocktail (1:1000), 200 mM AEBSF, and solubilization buffer overnight at 4 °C. The samples were centrifuged at 12000 rpm for 10 minutes and the supernatant was collected. 100 µL of an anti-HA magnetic bead solution (Invitrogen) were captured via a chilled magnetic stand and washed with 1X PBS/1x Protease Inhibitor Cocktail two times. The solubilized protein supernatant was added to the beads and incubated at 4 °C on a rotator overnight to allow binding of the native *Pf*ATAD3 complex to the anti-HA beads. The beads were collected using the magnetic stand and washed three times with Native Wash buffer (25 mM HEPES, 10 mM MgCl_2_, 50 mM KCl, 100 mM NaCl, 0.2% digitonin, 1x Protease Inhibitor Cocktail). The beads were then resuspended in Native Wash buffer/0.2% digitonin/1x Protease Inhibitor Cocktail and shipped to University of South Florida Mass spectrometry & Proteomic Core Facility for further processing and analysis. Data obtained from NF54 wildtype parasites was used as a negative control. LFQ intensity values were used for quantification and statistical analysis. Visualization of the data on a volcano plot was done on GraphPad Prism.

#### Transmission Electron Microscopy

NF54 *Pf*ATAD3-3xHA/TOM22-mNG parasites were synchronized with alanine (0.3 M in 10 mM HEPES, pH 7.6). Knockdown of *Pf*ATAD3 was induced at the ring stage by washout of aTc and the parasite culture was split into control and knockdown flasks. Late-stage parasites were then enriched using MACS Cell Separation Column (MiltenyiBiotec) at 36 hours and 84 hours after induction of *Pf*ATAD3 knockdown. The enriched parasites were then resuspended in their respective fresh culture media and maintained at proper culture conditions for about 1 hour to recover. The parasites were gently spun down at 100-300 ×*g* for 10 minutes in 15 mL conical-bottom polypropylene tubes. The pellet was resuspended in 3 mL of freshly prepared fixative (2% glutaraldehyde in 0.1 M Na-Cacodylate buffer), incubated for 5 minutes at room temperature, and kept at 4 °C. Further processing was performed by the Electron Microscopy Technician at Thomas Jefferson University, Philadelphia, PA including secondary fixation with 1% tannic acid, further staining with 2% OsO_4_ and 1% UA followed by agarose embedding, dehydration, and Spurr’s embedding. Thin sections were prepared on electron microscopy grids and imaged using a FEI Tecnai 12 120 keV digital Transmission Electron Microscope. Images were acquired at magnifications from 4kx to 42kx.

#### Bioinformatic Analyses

Amino acid sequences were retrieved using BLASTP software (https://00blast.ncbi.nlm.nih.gov/Blast.cgi?PAGE=Proteins) and protein family databases to find ATAD3A-like sequences from different phylogenetic groups. Multiple sequence alignment of the chosen FASTA complete sequences was done using MAFFT (Multiple Alignment using Fast Fourier Transform) on European Bioinformatics Institute online server (59). The feature annotations on the alignments were done using Jalview Software (60). Pairwise global alignment using BLOSUM62 as the scoring matrix was calculated between Human ATAD3 (UniProt ID: Q9NVI7) and *Pf*ATAD3 (UniProt ID: C0H4M1_PLAF7) and feature annotation was done using Jalview Software (60). The phylogenetic tree was inferred using maximum likelihood inference software accessed at NGPhylogeny.fr (61). Predictions of localization and essentiality (Mutagenesis Index Scores) of *Pf*ATAD3 and associated proteins were obtained from the *Plasmodium* bioinformatic database, PlasmoDB (62, 63). Predicted 3D structures were retrieved from the AlphaFold 3.0 Server (64).

#### Statistical Analyses

Unpaired t-tests were calculated for statistical significance of western blot and mtDNA copy number quantification on GraphPad Prism. Fold change was calculated using the median of the independent triplicate LFQ intensity raw values and a paired two-tailed distribution t-test analyses were conducted to determine statistical significance with a p-value cutoff of 0.05. 16-bit images were loaded onto FIJI and three lines of 50-pixel thickness each were drawn across the mitochondrial membranes to determine intensity measurements across the membrane. A region of interest within the mitochondrial matrix (lumen) was drawn to determine the intensity measurements within the matrix (lumen). Mean intensity measurements were subtracted from the maximum intensity measurements to calculate the inverted pixel values as the electron dense regions had lower pixel values. Calculated intensity measurements for the corresponding membrane were used to normalize the calculated lumen measurements, and an unpaired t-test was used to determine statistical significance on GraphPad Prism. To measure the width and corresponding area of mitochondrial TEM ultra sections, a 50-pixel thickness line was drawn across the smallest ends of the mitochondria. Unpaired t-test was used to calculate statistical significance on GraphPad Prism. Number of cells included for TEM statistical analyses: 36 h +*Pf*ATAD3, n = 11, 36 h -*Pf*ATAD3, n = 9, 84 h +*Pf*ATAD3, n = 12, 84 h -*Pf*ATAD3 = 12.

## Acknowledgments

We thank Hangjun Ke and Julie Verhoef for sharing the *Pf*TOM22-mNG and *Pf*Mito-mScarlet plasmids used in the generation of parasite lines in this study. We thank Gordon Ruthel at University of Pennsylvania PennVet Imaging Core for super-resolution laser scanning confocal imaging services, Dale Chaput at University of South Florida Mass Spectrometry and Proteomics Core Facility for proteomics services, and the Jefferson University Imaging Core for Transmission EM services.

