## Supplementary Information for "ATAD3 megadalton complex in *Plasmodium falciparum* is essential for mitochondrial and cellular viability"

### Figures

A

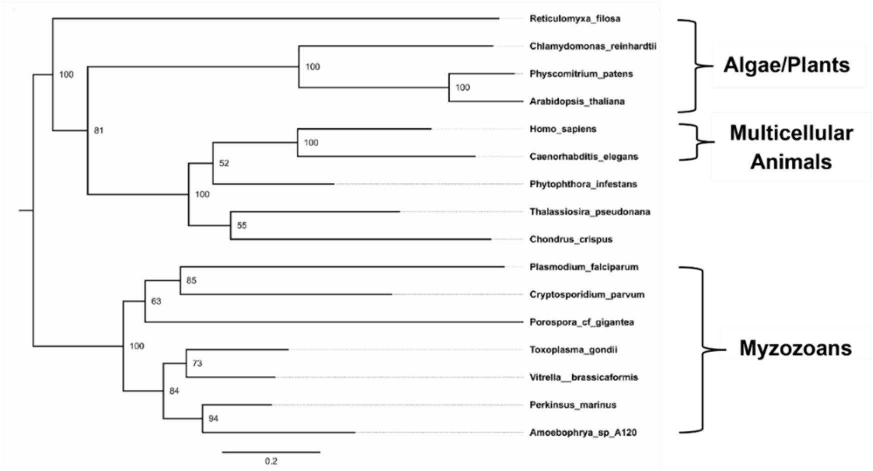

Reticulomyxa_filosa
Chlamydomonas_reinhardtii
Physcomitrium_patens
Arabidopsis_thaliana
Homo_sapiens
Caenorhabditis_elegans
Phytophthora_infestans
Thalassiosira_pseudonana
Chondrus_crispus
Plasmodium_falciparum
Cryptosporidium_parvum
Porospora_cf_gigantea
Toxoplasma_gondii
Vitrella__brassicaformis
Perkinsus_marinus
Amoebophrya_sp_A120
Algae/Plants
Multicellular Animals
Myzozoans
0.2


B

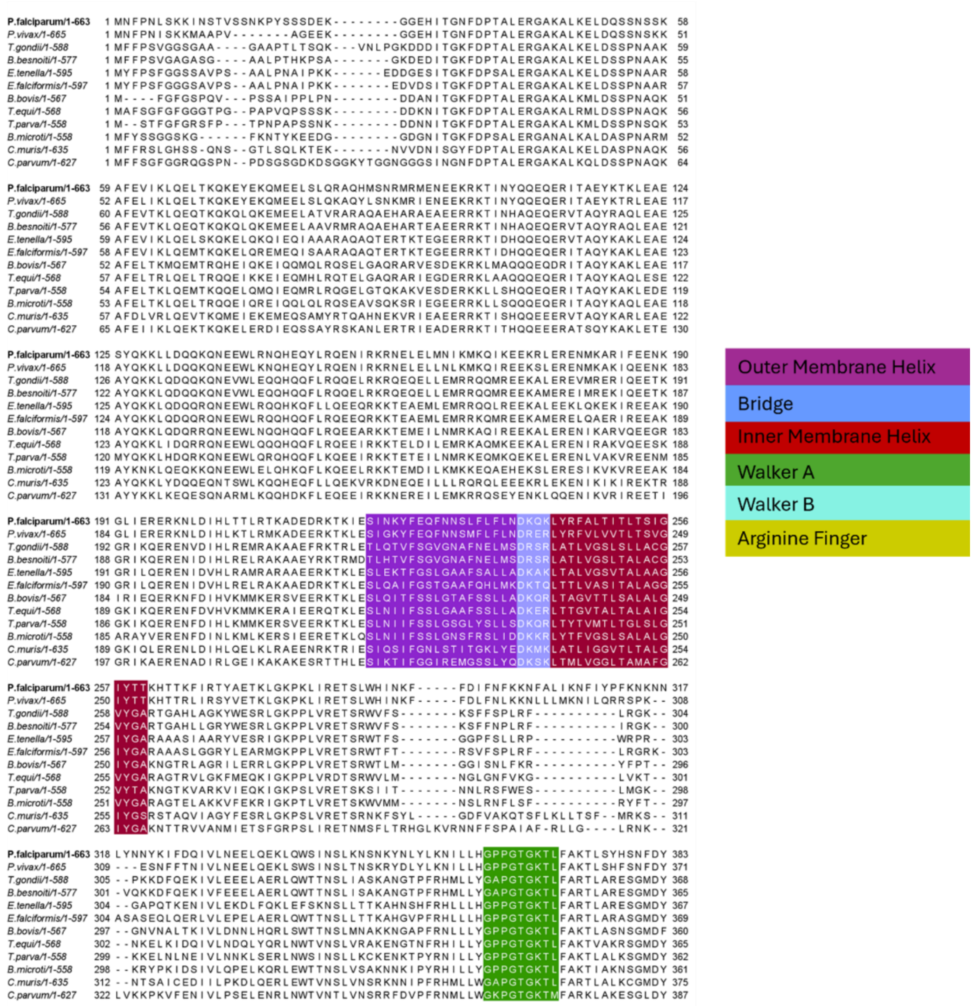

P.falciparum/1-663
P.vivax/1-665
T.gondii/1-588
B.besnoiti/1-577
E.tenella/1-595
E.falciformis/1-597
B.bovis/1-567
T.equi/1-568
T.parva/1-558
B.microti/1-558
C.muris/1-635
C.parvum/1-627
Outer Membrane Helix
Bridge
Inner Membrane Helix
Walker A
Walker B
Arginine Finger

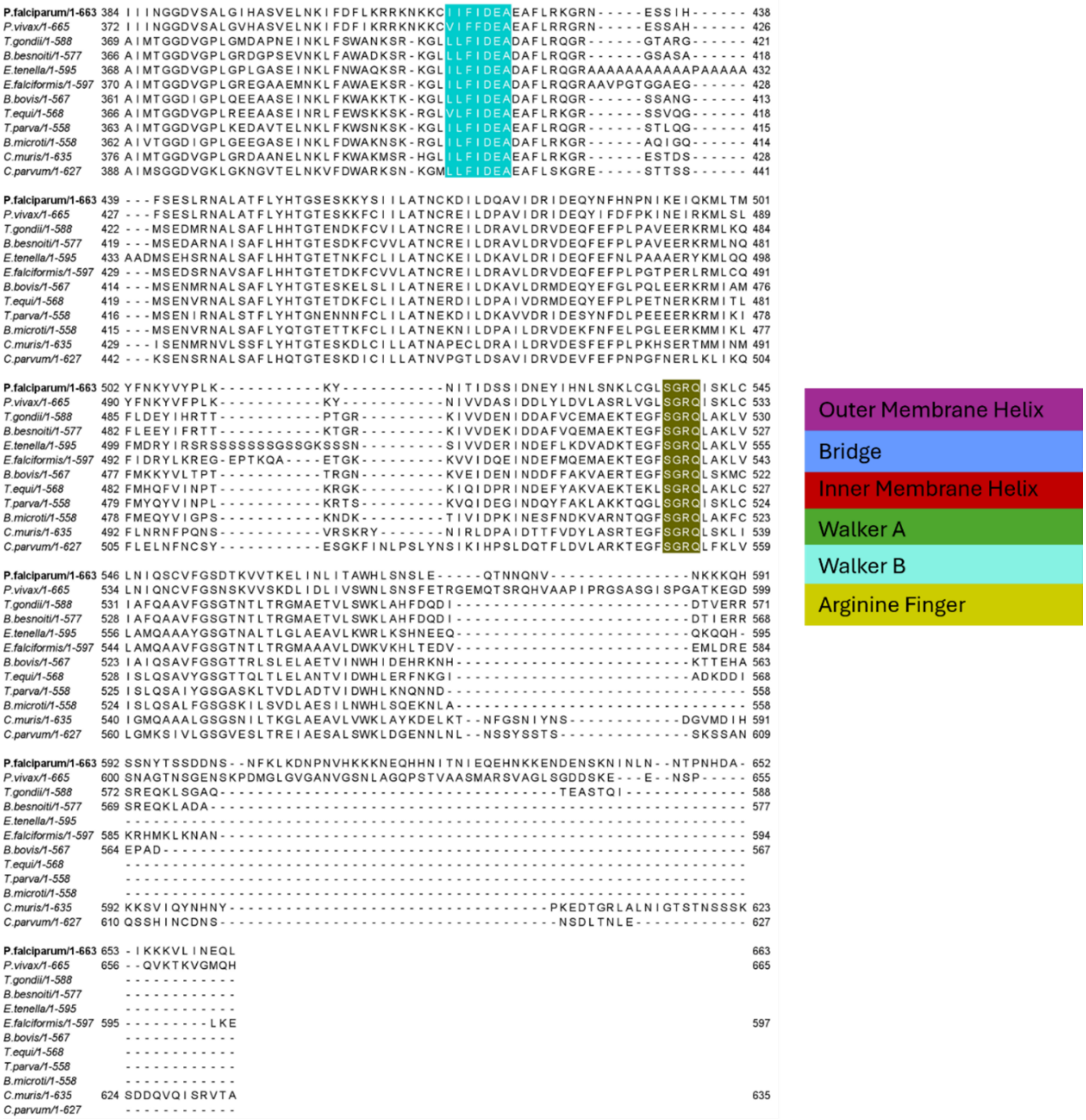


**S1. Fig (A)** Phylogenetic analysis showing the evolutionary relationship between orthologs of ATAD3 amongst alveolates and other eukaryotes including *Homo sapiens*, *Arabidopsis thaliana, Caenorhabditis elegans, Plasmodium falciparum, Toxoplasma gondii,* and *Cryptosporidium parvum*. **(B)** Multiple sequence analysis of ATAD3 orthologs in apicomplexan parasites.

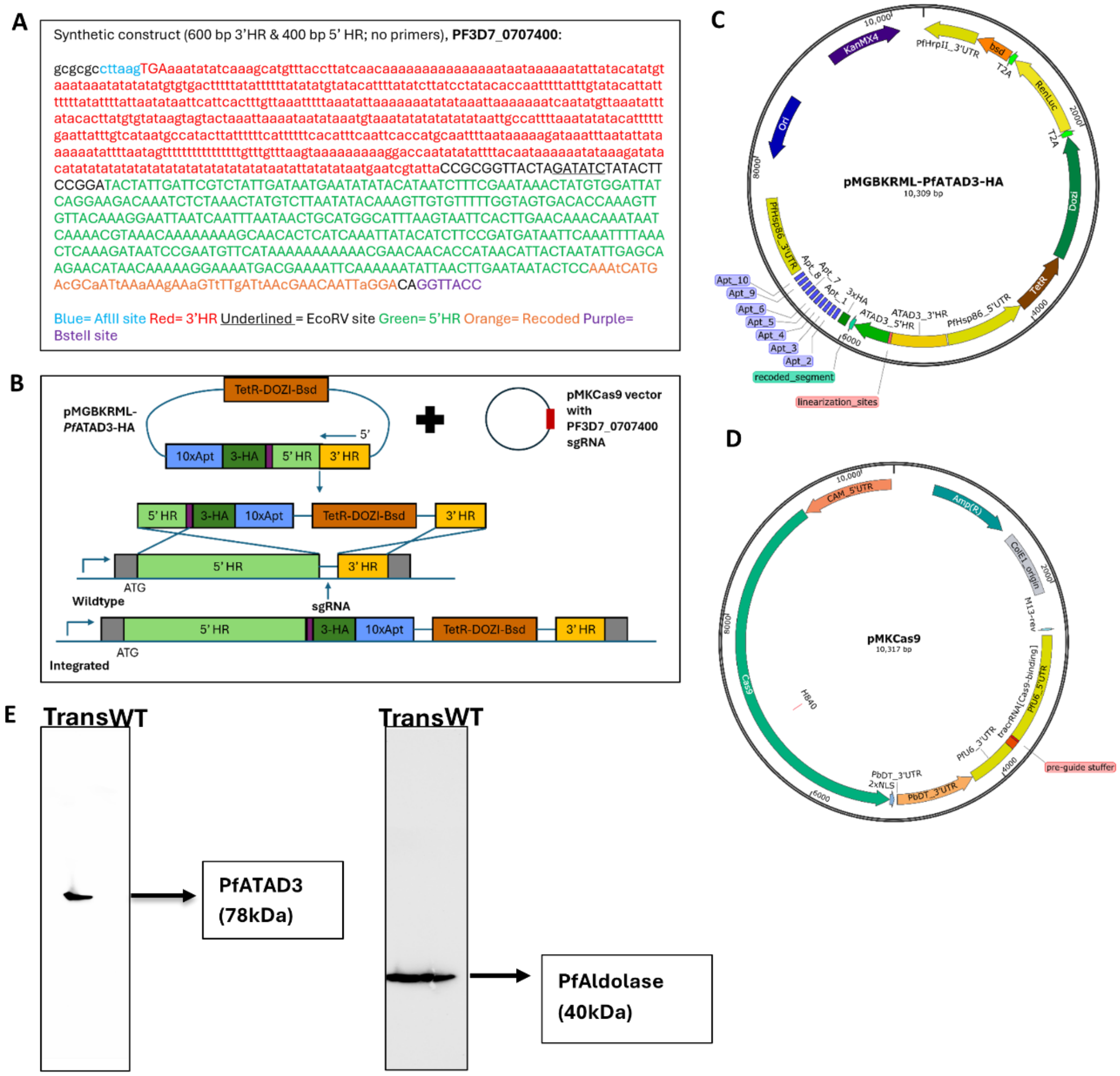


**S2. Fig (A)** Sequence of synthetic gene used to create the endogenously tagged *Pf*ATAD3A parasite line. The sequences are color coded and identified in a key below the synthetic construct. **(B)** CRISPR-Cas9 homologous recombination integration cloning strategies to generate endogenously tagged *Pf*ATAD3-HA-TetR, *Pf*ATAD3-HA-TetR/TOM22-mNG and *Pf*ATAD3-HA-TetR/mito-mScarlet transgenic parasite lines using corresponding parental lines. **(C)** Plasmid map of the pMGBKRML-*Pf*ATAD3-HA plasmid for endogenous tagging and conditional expression. Restriction sites and the identity of each region are indicated. Created with SnapGene. **(D)** Plasmid map of the pMKCas9 plasmid for guide RNA construction. Restriction sites and the identity of each region are indicated. Created with SnapGene. **(E)** Western Blot validating the *Pf*ATAD3-TetR/TOM22-mNG line using primary mouse anti-HA antibodies and primary rabbit anti-Aldolase antibodies.

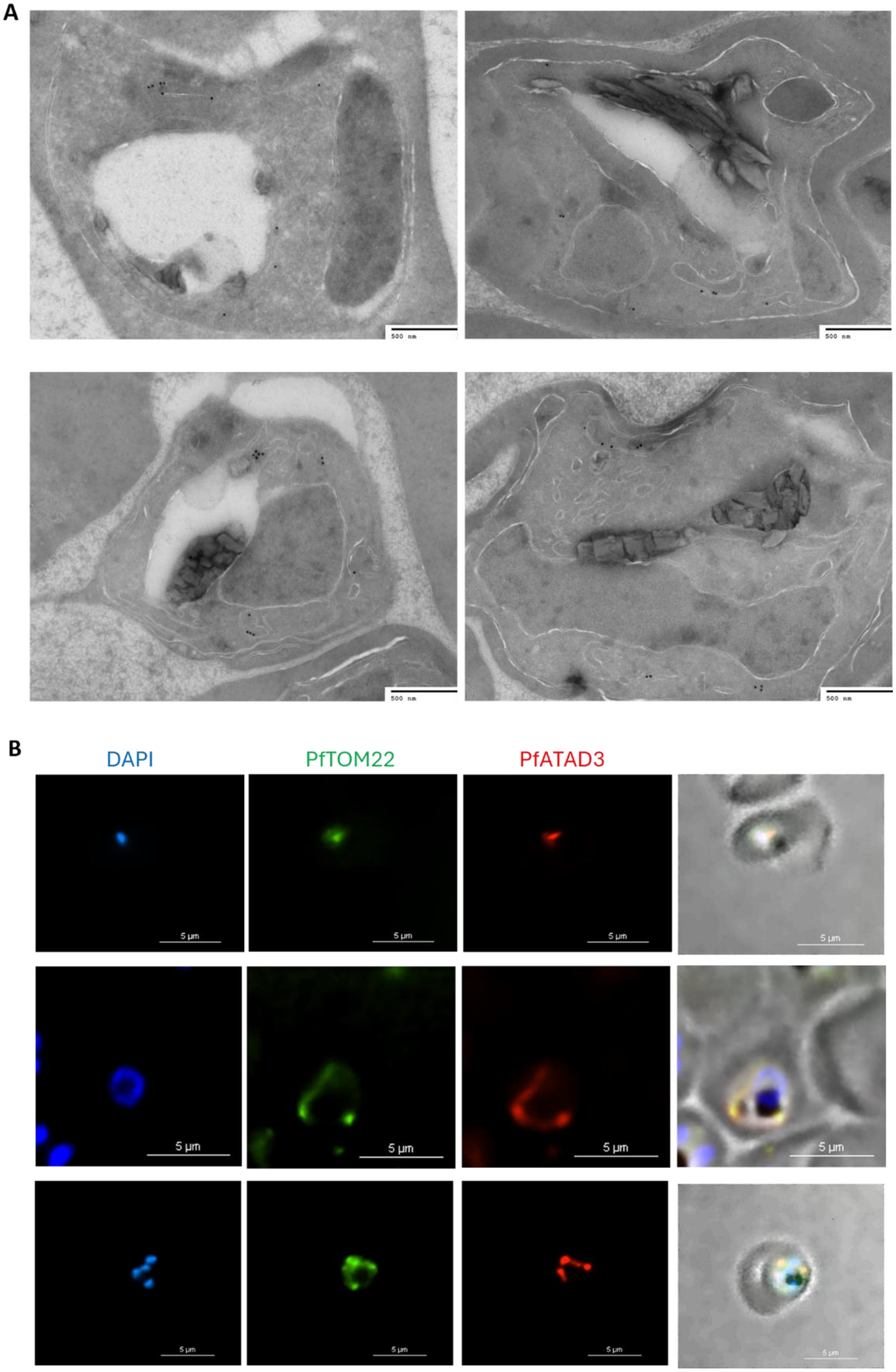


**S3. Fig (A)** Immuno-electron Micrographs showing localization of *Pf*ATAD3 to mitochondria of asexual parasites via mouse anti-HA or rabbit anti-HA primary antibodies and corresponding anti-mouse or anti-rabbit 18 nm colloidal gold particles. **(B)** Immunofluorescence Assay (IFA) demonstrating colocalization of *Pf*ATAD3 to *Pf*TOM22-mNeonGreen (Translocator of Outer Mitochondrial Membrane 22) in the mitochondria of asexual *P. falciparum* parasites.

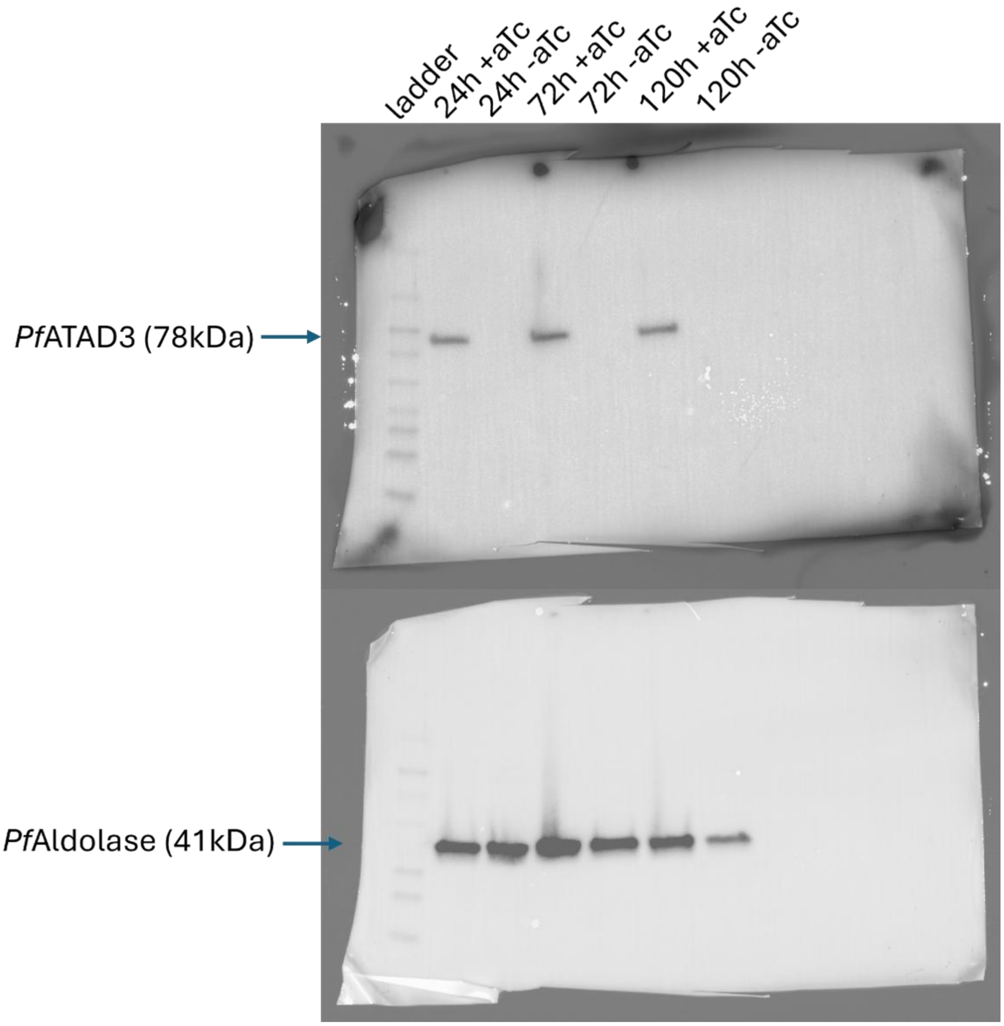


**S4 Fig.** Full gel image representative of the growth assay western blot.

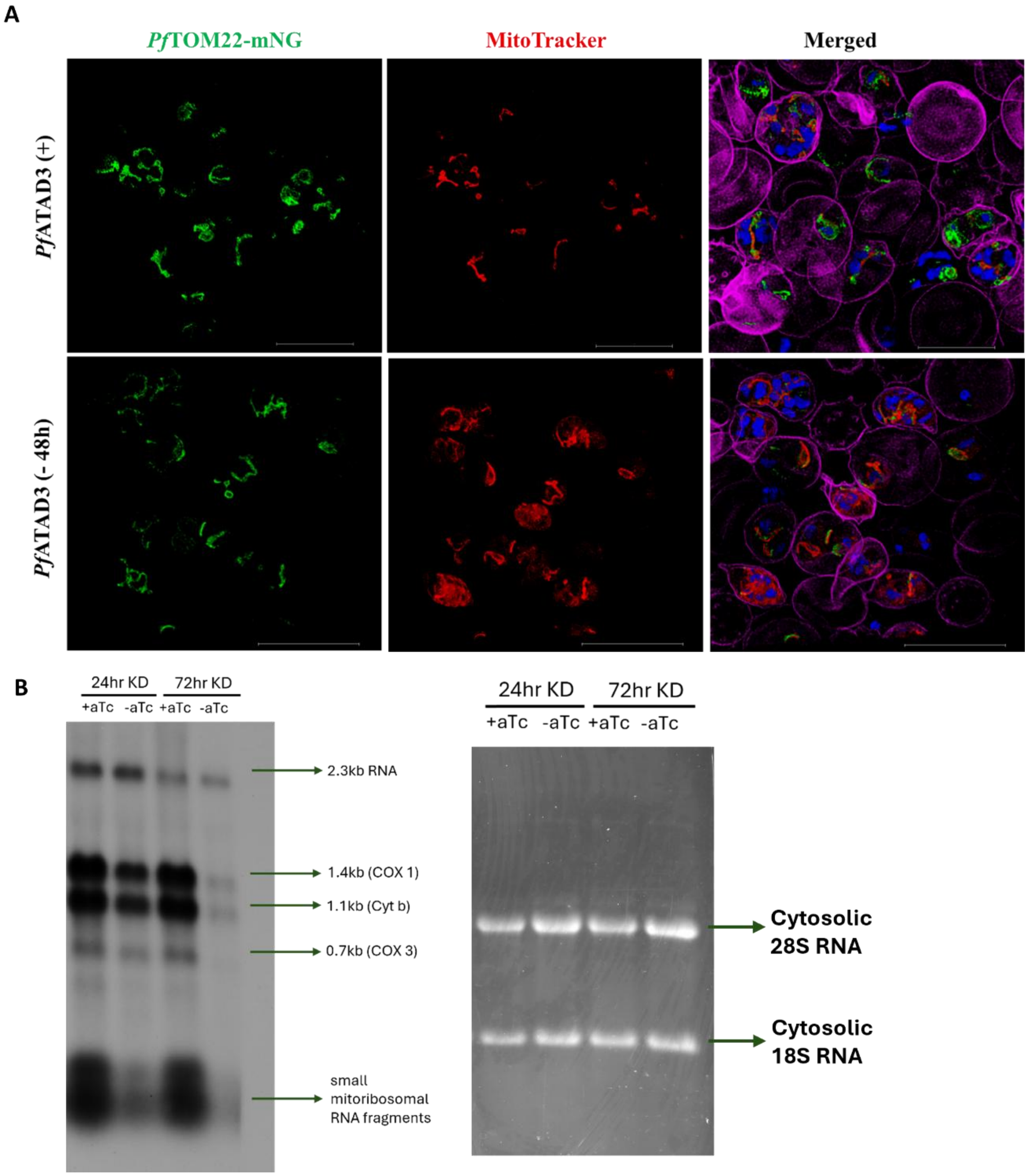


**S5 Fig. (A)** Super-resolution live cell imaging of mitochondrial membrane potential upon loss of *Pf*ATAD3. Partial loss of mitochondrial membrane potential observed during first asexual life cycle (48 h) post *Pf*ATAD3 knockdown. *Pf*DNA – DAPI (Blue); Outer mitochondrial membrane – *Pf*TOM22-mNG (Green); Mitochondrion – MitoTracker (Red); Host red blood cell – Wheat Germ Agglutinin (Purple). Scale bar is 10 µm. **(B)** Full northern blot (left) and denaturing agarose RNA gel (right) of one representative replicate demonstrating reduction of processed mitochondrial RNA transcripts upon knockdown of *Pf*ATAD3.

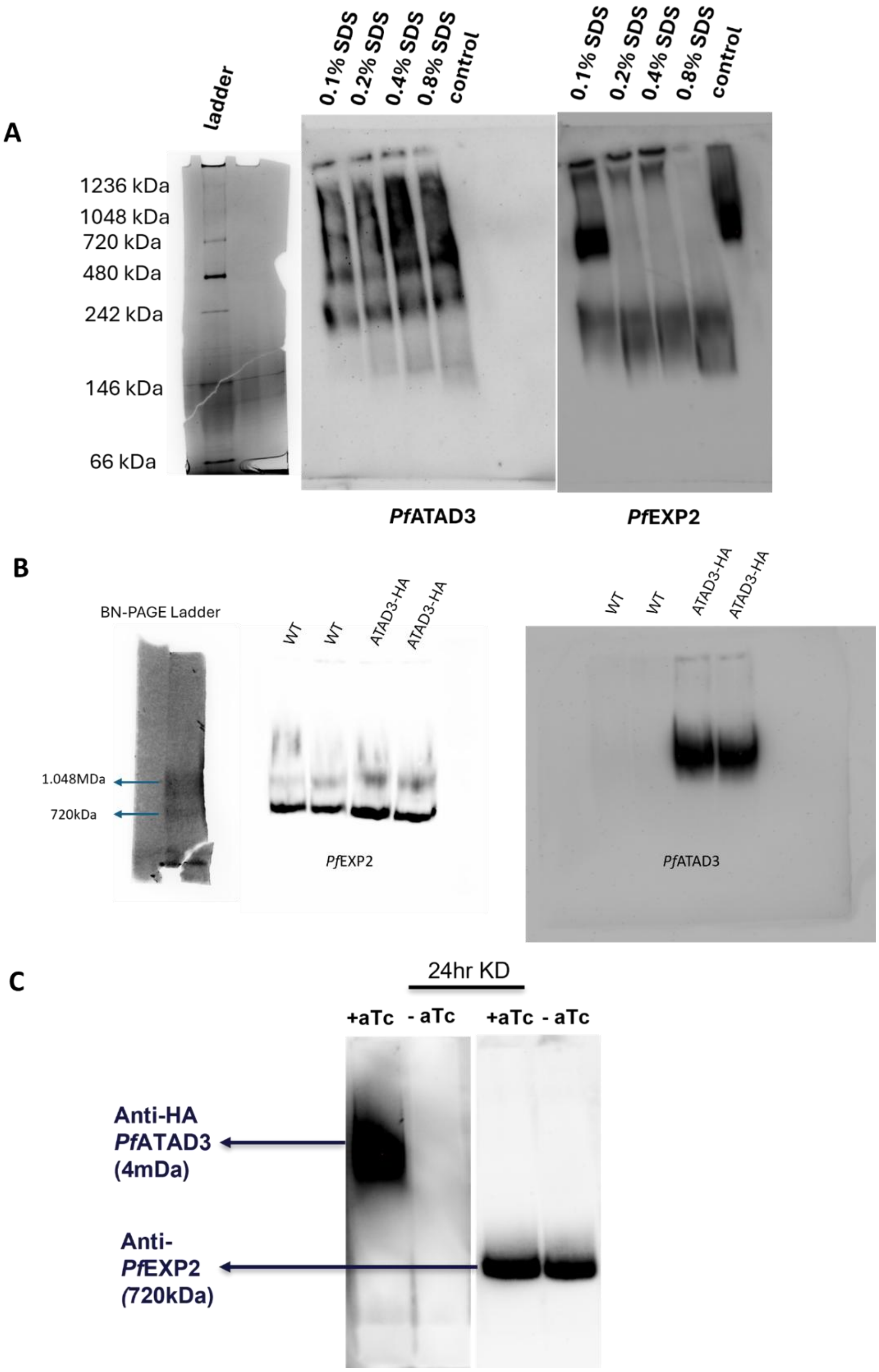


**S6 Fig. (A)** Blue-Native PAGE of control and SDS-treated solubilized extract demonstrating the presence of multiple sub-complexes of *Pf*ATAD3. *Pf*EXP2 was used as a loading control. **(B)** Full large pore composite native gel showing *Pf*ATAD3 is present in a mega-Dalton hetero-oligomeric complex. **(C)** Large pore composite native gel showing aTc washout for 24h induces knockdown of the giant megaDalton *Pf*ATAD3 complex.

**S1 Table. Enriched Interacting Protein Partners of *Pf*ATAD3-HA in asexual *P. falciparum* parasites.**

| Gene | Protein Name | Fold Change | P-value | MIS | Prior Localization Information |
|---|---|---|---|---|---|
| PF3D7_1102700 | early transcribed membrane protein 11.1 | 9.43E+08 | 0.018038 | 1 | Symbiont-containing vacuole membrane |
| PF3D7_0707400 | ATPase family AAA domain-containing protein 3A, putative | 7.38E+08 | 0.010399 | 0.13 | Mitochondrion |
| PF3D7_1469200 | shewanella-like protein phosphatase 1, putative | 72738111 | 0.034519 | 0.162 | Nucleus, Cytoplasm, Endoplasmic Reticulum |
| PF3D7_1330600 | elongation factor Tu, putative | 51627111 | 0.000307 | 0.131 | Mitochondrion, Plastid, Nucleus |
| PF3D7_1119600 | ATP-dependent zinc metalloprotease FTSH, putative | 48661111 | 0.008991 | 0.141 | Mitochondrial Inner Membrane |
| PF3D7_0726900 | mitochondrial import inner membrane translocase, TIM50 | 43172111 | 0.009813 | 0.12 | Mitochondrial Inner Membrane |
| PF3D7_1430200 | plasmepsin IX | 42511111 | 0.000571 | 0.238 | Rhoptry |
| PF3D7_1356200 | mitochondrial import inner membrane translocase, TIM23 | 38442111 | 0.013472 | 0.12 | Mitochondrial Inner Membrane |
| PF3D7_0212900 | arginyl-tRNA--protein transferase | 34974111 | 0.032805 | 0.999 | Cytoplasm |
| PF3D7_0413500 | phosphoglucomutase-2 | 34916111 | 0.027304 | 0.119 | Cytoplasm, Mitochondrion Membrane |
| PF3D7_0419600 | ran-specific GTPase-activating protein 1, putative | 34629111 | 0.045771 | 0.998 | Cytoplasm, Nuclear Pore, Nucleus |
| PF3D7_1468100 | MORC family protein | 31152111 | 0.012493 | 0.179 | Nucleus |
| PF3D7_1462300 | GTP-binding protein, putative | 29693111 | 0.013979 | 0.178 | Nucleus |
| PF3D7_1104100 | syntaxin, Qa-SNARE family | 24921111 | 0.009559 | 0.643 | Vesicle Membrane, Food vacuole, Plasma Membrane |
| PF3D7_0303000 | N-ethylmaleimide-sensitive fusion protein | 16594111 | 0.020639 | 0.186 | Golgi stack, Food vacuole, Host cell cytoplasm |
| PF3D7_0408500 | flap endonuclease 1 | 15149111 | 0.004239 | 0.999 | Mitochondrion, Nucleolus, Nucleoplasm |
| PF3D7_0207500 | serine repeat antigen 6 | 8528411 | 0.010314 | 0.252 | Symbiont-containing vacuole membrane |
| PF3D7_0320800 | ATP-dependent RNA helicase DDX6 | 19.19762 | 0.004638 | 1 | P-body, Cytoplasm, Cytoplasmic stress granule, Nucleus |

| Gene | Protein Name | Fold Change | P-value | MIS | Prior Localization Information |
|---|---|---|---|---|---|
| PF3D7_1434800 | mitochondrial acidic protein MAM33, putative | 11.78068 | 0.017136 | 0.509 | Mitochondrial Matrix |
| PF3D7_0617900 | histone H3 variant | 8.421096 | 0.041978 | 0.411 | Nucleosome |
| PF3D7_0610400 | histone H3 | 5.103586 | 0.040886 | 0.176 | Nucleosome |
| PF3D7_1441100 | conserved Plasmodium protein, unknown function | 4.188605 | 0.04055 | 1 | Nucleus |

**S2 Table. Primer sequences used in parasite line generation**

| Primer Name | Sequence |
|---|---|
| 0707400FWD | GATGTACCTAGGtaaaATGAATTTTCCTAATTTGAGTAAGAAAATAAATTC |
| 0707400REV | GATGTACGTACGgttCAATTGTTCATTTATTAATACCTTTTTCTTTATAGC |
| PfTom22 FWD | cgTGTACAAGggatctggatctCGTACGATGGGAACAGCACTATCAAAAATTATTACG |
| PfTom22 REV | gtCTTAAGTTAGTTTAATTGTGGAACATTGGC |
| PF3D7_0707400 gRNA1 | CATATTAAGTATATAATATTGTTCTTTATAGCATCATGGTTGTTTCAGAGCTATGCTGGA |
| PF3D7_0707400 gRNA2 | atttCATATTAAGTATATAATATTGCCATGATGCTATAAAGAAAAGTTTCAGAGCTATGCTGGAaac |

**S3 Table. Digital PCR Custom Designed TaqMan Gene Expression Assay Details**

| Assay ID | Assay Name | Assay Mix Concentration | Reporter 1 Dye | Reporter 1 Concentration (µM) | Reporter 1 Quencher | Forward Primer Concentration (µM) | Reverse Primer Concentration (µM) | Forward Primer Sequence | Reverse Primer Sequence | Reporter 1 Sequence | Context Sequence |
|---|---|---|---|---|---|---|---|---|---|---|---|
| AP33C2U | CYTOCHROME_B | 20x | FAM | 5 | NFQ | 18 | 18 | TGTACTACATTTTATCTTACCATTTATTGGATTATGTATTGT | GGGTATTTTTAATGCTGTATCATACCCT | CATGGTAGCACAAATC | TGTACTACATTTTATCTTACCATTT |
| AP7DY7N | GAPDH | 20x | VIC | 5 | NFQ | 18 | 18 | TGGTCAATTTCCATGTGAGGTAACC | GATCCTTTTCAGCAAAAACACTGACT | CACGCTGATGGATTTT | TGGTCAATTTCCATGTGAGGTAACC |

**S1 Movie.** 3D super-resolution live cell imaging videos demonstrating mitochondrial membrane potential in *Pf*ATAD3 (+) parasites. *Pf*DNA – DAPI (Blue); Outer mitochondrial membrane – *Pf*TOM22-mNG (Green); Mitochondrion – MitoTracker (Red); Host red blood cell – Wheat Germ Agglutinin (Purple). Scale bar is 10 µm.

**S2 Movie.** 3D super-resolution live cell imaging videos demonstrating mitochondrial membrane potential in *Pf*ATAD3(-) parasites. *Pf*DNA – DAPI (Blue); Outer mitochondrial membrane – *Pf*TOM22-mNG (Green); Mitochondrion – MitoTracker (Red); Host red blood cell – Wheat Germ Agglutinin (Purple). Scale bar is 10 µm.
